# Fronto-Parietal Oscillatory Dynamics of Emotion Regulation as a Function of Adult Attachment Orientations

**DOI:** 10.1101/2025.11.22.689931

**Authors:** Marcos Domic-Siede, Javiera Figueroa-Cuevas, Krishna Leiva-Cortés, Daniela López, Mónica Guzmán-González, Thomas Lau-Lemus, Sara Hernández, Romina Ortiz, Lesly González, Jaime R. Silva

**Author notes:** Corresponding Author: Marcos Domic-Siede. Correspondence concerning this article should be addressed to Marcos Domic-Siede, Laboratorio de Neurociencia Cognitiva, Escuela de Psicología, Universidad Católica del Norte, Av. Angamos 0610, Antofagasta, Chile.

## Abstract

Emotion regulation enables individuals to modulate emotional experiences and behaviors according to situational demands. Within attachment theory, individual differences in attachment anxiety and avoidance are conceived as interpersonal dispositions that influence the quality and efficiency of emotion regulation strategies, potentially shaping the underlying neural dynamics. This study examined cortical oscillatory activity during two regulation strategies—cognitive reappraisal and expressive suppression—focusing on theta (4–8 Hz) and beta (15–30 Hz) bands at the source level. Forty adults (21 male, 18 female, 1 gender-unspecified; *M* = 27.58 years, *SD* = 8.71) performed an emotion regulation task involving emotionally evocative images from the International Affective Picture System (IAPS) while EEG was recorded. Cortical sources were reconstructed using standardized low-resolution brain electromagnetic tomography (sLORETA). Linear mixed-effects models (LMMs) assessed the effects of condition, region of interest (ROI), and attachment orientations on oscillatory power. Results showed that higher attachment anxiety predicted reduced theta power in the right dorsolateral prefrontal cortex (dlPFC) during reappraisal, indicating attenuated recruitment of cognitive control mechanisms. In the beta band, suppression reduced activity in the right parietal lobe/precuneus — a region involved in self-referential and attentional processes — across participants, while during reappraisal, attachment anxiety was linked to lower beta power and attachment avoidance to higher beta power in the left dlPFC. Together, these findings indicate that cognitive reappraisal and expressive suppression engage distinct cortical oscillatory systems, and that interpersonal dispositions modulate their neural implementation. Frontal theta appears to index top-down control during reappraisal, whereas beta activity shows a dual pattern: frontal beta variations reflect differences in control stability associated with attachment orientations, and parietal beta decreases during suppression suggest attentional disengagement from self-referential processing.

## 1. Introduction

### 1.1 Attachment Theory

Attachment theory describes an innate motivational system that promotes proximity-seeking behavior toward a caregiver under threat, providing safety and emotional regulation (Bowlby, 1969, 1973, 1982; Mikulincer & Shaver, 2016). Through early interactions, individuals develop internal working models of the self and others that shape expectations about relationships (Bowlby, 1973; Brennan et al., 1998). In adulthood, attachment is commonly conceptualized along two dimensions: anxiety and avoidance (Brennan et al., 1998; Mikulincer & Shaver, 2016; Guzmán-González et al., 2020). Attachment anxiety involves a negative view of the self and a fear of rejection or abandonment, leading to hyperactivating strategies such as clinging and heightened emotional expression. Attachment avoidance reflects a negative view of others and discomfort with intimacy, promoting deactivating strategies such as emotional suppression and withdrawal (Mikulincer & Shaver, 2003; Bartholomew & Horowitz, 1991; Hazan & Shaver, 1987). Secure attachment (low anxiety and avoidance) is characterized by positive expectations about self and others, supporting more adaptive emotional responses (Mikulincer & Shaver, 2016; Messina et al., 2024). In contrast, attachment insecurity—whether anxious or avoidant—is consistently linked to emotion regulation difficulties and maladaptive affective responses (Cassidy & Shaver, 2016; Guzmán-González et al., 2016; Han & Kahn, 2017; Henschel et al., 2020; Long et al., 2020; Vrtička & Vuilleumier, 2012).

### 1.2. Emotion Regulation

Emotion regulation refers to the processes through which individuals influence which emotions they experience, when they occur, and how they are expressed (Gross, 1998, 2002). In Gross’s process model (Gross, 1998; Olderbak et al., 2023), two key families of strategies are cognitive change and response modulation. Cognitive change includes cognitive reappraisal, which involves reframing an emotion-eliciting situation to alter its emotional impact (Gross, 2002; Uusberg et al., 2019). Response modulation includes expressive suppression, or the conscious inhibition of emotional expressions (Gross, 2002; Goldin et al., 2008; Olderbak et al., 2023). Not all strategies are equally adaptive: reappraisal is generally linked to higher well-being, lower physiological cost, and better social functioning, while suppression is associated with increased distress and poorer relational outcomes (Gross & John, 2003; Goldin et al., 2008; Olderbak et al., 2023).

Attachment orientations strongly shape both the selection and effectiveness of regulation strategies. Individuals with higher attachment security tend to use reappraisal more effectively and flexibly, whereas attachment anxiety and avoidance are associated with maladaptive regulation patterns (Mikulincer & Shaver, 2016; Guzmán-González et al., 2016; Long et al., 2020; Ramos-Henderson et al., 2024; Domic-Siede et al., 2024; Vrtička & Vuilleumier, 2012). Anxiously attached individuals often show emotional hyperactivation, rumination, and reduced control (Henschel et al., 2020; Ramos-Henderson et al., 2024), while avoidantly attached individuals tend to rely on suppression and show reduced efficacy in reappraisal (Vrtička & Vuilleumier, 2012).

Emotion regulation is not only an individual cognitive process but also a socially embedded one. Strategies such as reappraisal and suppression are shaped by early relational experiences and expectations about others’ availability and responsiveness (Mikulincer & Shaver, 2016). From this perspective, the ability to regulate emotion develops in the context of interpersonal relationships, particularly attachment bonds, which guide how individuals manage affect in social contexts throughout life.

### 1.3. Neural Bases of Emotion Regulation

Emotion regulation is implemented by a set of interconnected neural systems involving prefrontal, limbic, and parietal regions. Neuroimaging studies, primarily using functional magnetic resonance imaging (fMRI), have consistently implicated the prefrontal cortex (PFC) as a critical region for top-down regulation of emotional responses, particularly in down-regulating activity in emotion-generative structures like the amygdala (Buhle et al., 2014; Morawetz et al., 2017; Ochsner et al., 2002, 2012). Within the PFC, different subregions have specialized functions. The dorsolateral prefrontal cortex (dlPFC) is central for cognitive control and working memory, supporting the manipulation of information required for reappraisal (Buhle et al., 2014). The ventrolateral prefrontal cortex (vlPFC) contributes to the selection and inhibition of responses, particularly in relation to suppressing inappropriate emotional behaviors (Morawetz et al., 2017). The dorsomedial prefrontal cortex (dmPFC) and anterior cingulate cortex (ACC) are implicated in conflict monitoring and self-referential processing, playing a role in evaluating the success of regulatory efforts (Etkin et al., 2015; Ochsner et al., 2012). Finally, parietal regions, including the inferior parietal lobule and precuneus, contribute to attentional reorienting and visuospatial processing during regulation. According to classic cortical organization models (Northoff et al., 2006), the precuneus—part of the medial cortical system—also supports self-referential processing, linking internal representations of the self with ongoing emotional and cognitive states (Ochsner et al., 2012).

The amygdala, a key limbic structure, is reliably activated during emotional experience and is commonly targeted by regulation processes. Successful reappraisal is associated with decreased amygdala activation, which reflects reduced emotional reactivity (Buhle et al., 2014; Goldin et al., 2008). Conversely, when individuals engage in suppression or when regulation fails, amygdala activity often remains high or increases (Ochsner et al., 2012). These findings support the idea that reappraisal is more efficient than suppression in terms of attenuating the neural correlates of negative affect.

The PFC–amygdala interaction thus forms a core circuit for regulating emotion: PFC regions implement cognitive control strategies that down-regulate amygdala responses to emotional stimuli (Etkin et al., 2015). Connectivity studies suggest that stronger inverse coupling between the PFC and amygdala (i.e., greater prefrontal control over limbic reactivity) predicts better regulation outcomes and lower affective symptoms (Morawetz et al., 2017; Ochsner et al., 2012).

This frontolimbic model has informed much of the contemporary understanding of emotion regulation at the neural level.

Critically, evidence linking attachment orientations with frontolimbic circuitry remains mixed. While several studies have reported that securely attached individuals show more efficient prefrontal down-regulation of amygdala activity during emotional challenges (Vrtička et al., 2008, 2012; Vrtička & Vuilleumier, 2012), findings across the literature are heterogeneous. A recent meta-analysis suggests that the most consistent neural differences involve the lateral prefrontal cortex, typically showing reduced activation among avoidantly attached individuals, and the amygdala, which tends to exhibit hyperactivation in anxiously attached individuals (Ran & Zhang, 2018). However, these patterns vary considerably across paradigms and task demands, as highlighted by more recent reviews (Long et al., 2020). Anxious attachment has been linked to heightened limbic responsivity and inefficient prefrontal recruitment, reflecting increased sensitivity to social threat and difficulty down-regulating distress, whereas avoidant attachment often shows blunted amygdala reactivity together with attenuated or disengaged lateral prefrontal activity, consistent with deactivating or suppression-based strategies. Overall, the data point to context-dependent modulations within frontolimbic networks, emphasizing variability in how attachment dispositions shape emotional control at the neural level.

Moreover, electrophysiological measures have begun to complement fMRI findings by offering fine-grained temporal resolution of emotion regulation processes. Event-related potentials (ERPs) such as the late positive potential (LPP) have been used to index sustained attentional engagement with emotional stimuli, and its modulation during reappraisal or suppression provides insight into the time course of regulatory effort (Hajcak & Nieuwenhuis, 2006; Moser et al., 2006). ERP studies have found that attachment anxiety is associated with increased LPP amplitude to negative stimuli, consistent with emotional hyperreactivity (Ramos-Henderson et al., 2024; Domic-Siede et al., 2025a).

Together, fMRI and EEG studies indicate that the neural bases of emotion regulation are deeply intertwined with attachment orientations, influencing both the effectiveness and preferred strategies of emotional control.

### 1.4 Brain Oscillations

Electroencephalography (EEG) provides a powerful tool for examining the temporal dynamics of neural processes underlying emotion regulation. EEG oscillatory activity—i.e., rhythmic patterns in neuronal firing—reflects different modes of neural communication and support cognitive and affective functions (Başar et al., 2001; Buzsáki & Draguhn, 2004; Klimesch, 1999). Among the various frequency bands, theta (4–8 Hz) and beta (13–30 Hz) oscillations have been most consistently implicated in emotion regulation (Abid et al., 2025; Domic-Siede et al., 2024, 2025b; Knyazev, 2007; Peng et al., 2025). Frontal midline theta activity is commonly associated with cognitive control and mental effort, especially in tasks requiring conflict resolution, working memory, or sustained attention (Cavanagh & Frank, 2014). During emotion regulation tasks, increases in theta power have been interpreted as reflecting the engagement of top-down regulatory processes, such as reappraisal (Ertl et al., 2013). Theta synchronization is believed to support the functional integration of distributed cortical and subcortical systems (Cavanagh & Frank, 2014), facilitating the implementation of regulatory strategies in real time. Beta activity, by contrast, has been associated with both motor inhibition and the maintenance of cognitive and emotional states (Engel & Fries, 2010; Heinrichs-Graham & Wilson, 2016; Gilbertson et al., 2005). In the context of emotion regulation, beta oscillations—particularly over frontal and parietal regions—are thought to reflect the effortful inhibition of emotional expression (Davidson et al., 2000; Domic-Siede et al., 2025b). Thus, theta and beta oscillations provide complementary indices of regulatory engagement: theta signaling controlled modulation of emotion, and beta reflecting the inhibition of emotional expression.

Importantly, individual differences in attachment orientations may influence the pattern of oscillatory activity during regulation. Preliminary findings suggest that individuals higher in attachment anxiety exhibit decreased frontal theta activity and a reduced frontal theta synchrony during emotional tasks, consistent with increased regulatory effort or emotional reactivity (Domic-Siede et al., 2024, 2025b). Individuals higher in attachment avoidance, in turn, may show altered beta patterns, possibly reflecting habitual suppression or disengagement from emotional stimuli (Domic-Siede et al., 2025b). These findings offer a promising avenue for linking attachment theory to the neurophysiology of emotion regulation, with oscillatory markers providing real-time indicators of how regulatory processes unfold across individuals.

### 1.5 The Present Study

Building on the theoretical and empirical frameworks reviewed above, the present study aimed to investigate the cortical oscillatory dynamics underlying two emotion regulation strategies—cognitive reappraisal and expressive suppression—and their modulation by attachment orientations. Specifically, we focused on theta (4–8 Hz) and beta (15–30 Hz) frequency bands at the source level to explore the neural mechanisms associated with cognitive control and expressive suppression, respectively. The integration of attachment theory into the study of oscillatory brain activity represents an emerging direction in affective neuroscience—one that bridges individual differences in socio-emotional motivation with neurophysiological mechanisms of regulation (Long et al., 2020; Vrtička & Vuilleumier, 2012).

Individuals with higher attachment anxiety often exhibit hyperactivating responses—heightened emotional reactivity and difficulties engaging prefrontal control—whereas those with higher attachment avoidance tend to engage in deactivating responses, such as emotional suppression and withdrawal from affective cues (Long et al., 2020; White et al., 2020). These tendencies suggest distinct neural signatures that may be observable in oscillatory activity.

At the neural level, we expected frontal theta activity to index top-down control during cognitive reappraisal (Cavanagh & Frank, 2014; Ertl et al., 2013) and beta oscillations to reflect inhibitory control during expressive suppression (Engel & Fries, 2010; Heinrichs-Graham & Wilson, 2016). We therefore hypothesized that:

1. Higher **attachment anxiety** would predict reduced frontal theta during reappraisal, indicating less efficient prefrontal engagement.
2. Higher **attachment avoidance** would predict increased beta power during suppression, consistent with enhanced motor inhibition and emotional disengagement.

To test these hypotheses, we used source-level EEG analyses based on a multi-atlas anatomical framework (Desikan et al., 2006; Destrieux et al., 2010; Klein & Tourville, 2012). This approach allowed us to estimate oscillatory activity within well-defined cortical regions involved in emotion regulation, including the anterior cingulate cortex (ACC), medial and dorsolateral prefrontal cortices (mPFC, dlPFC), ventrolateral prefrontal cortex/inferior frontal gyrus (vlPFC/IFG), supplementary and pre-supplementary motor areas (SMA/pre-SMA; BA 6), and parietal cortex/precuneus (PL/Precuneus).

By combining attachment theory with electrophysiological evidence, this study sought to elucidate how interpersonal motivational systems modulate neural oscillatory mechanisms of emotion regulation. Such findings may contribute to an integrative understanding of the social and neurophysiological architecture of emotional control, offering novel insights into the interplay between attachment, cognition, and affect at the cortical level.

## 2. Materials and Methods

### 2.1. Participants

A purposive, non-probabilistic sampling strategy was used to recruit adult participants (≥18 years old) with normal or corrected-to-normal vision and no history of severe neuropsychiatric disorders. All participants provided written informed consent prior to participation. Recruitment was conducted through digital platforms of the School of Psychology at Universidad Católica del Norte, Chile.

The initial sample comprised 46 participants. Six were excluded due to excessive EEG noise or technical issues, resulting in a final sample of 40 adults (38 Chilean, 1 Argentinian, and 1 Bolivian): 18 females (M = 27.44 years, SD = 8.33; range = 19–52), 21 males (M = 27.81 years, SD = 9.12; range = 19–57), and one participant with non-specified sex/gender (age = 26). The overall sample had a mean age of M = 27.58 years (SD = 8.71; range = 19–57) (**Table 1**).

**Table 1.** Sociodemographic characteristics of the participants (n = 40)

| Variables | n | % | Mean (SD) |
| --- | --- | --- | --- |
| **Age** |  |  | 27.58 (8.60) |
| **Sex** |  |  |  |
| Male | 21 | 52.50 |  |
| Female | 18 | 45.00 |  |
| Other | 1 | 2.50 |  |
| **Nationality** |  |  |  |
| Chilean | 38 | 95.00 |  |
| Bolivian | 1 | 2.50 |  |
| Argentine | 1 | 2.50 |  |
| **Marital status** |  |  |  |
| Single | 33 | 82.50 |  |
| Married | 7 | 17.50 |  |
| **Socioeconomic level** |  |  |  |
| Low | 1 | 2.50 |  |
| Low–Middle | 21 | 52.50 |  |
| Upper–Middle | 18 | 45.00 |  |
| High | 0 | 0.00 |  |
| **Education** |  |  |  |
| Completed Secondary Education | 3 | 7.50 |  |
| Completed Technical Education | 4 | 10.00 |  |
| Completed Higher Education | 9 | 22.50 |  |
| Incomplete Technical/Higher Education | 20 | 50.00 |  |
| Postgraduate Studies | 4 | 10.00 |  |
| **Occupation** |  |  |  |
| Employed | 20 | 50.00 |  |
| Homemaker | 2 | 5.00 |  |
| Technical/Higher Education Student | 18 | 45.00 |  |

An *a priori* power analysis was conducted using G*Power 3.1.9.2 (Faul et al., 2007) for a repeated-measures ANOVA with three within-subject conditions (α = .05, power = .85, assumed correlation among measures = .50, nonsphericity correction = 1). The analysis indicated that a minimum of 31 participants was required to detect a medium effect size (*f* = 0.25). The final sample of 40 participants exceeded this threshold, ensuring sufficient statistical power for detecting effects of interest. To assess sensitivity, a post-hoc power analysis considered the significant interactions from our linear mixed-effects models: (i) attachment anxiety × reappraisal in the theta band at right dlPFC (β ≈ −0.19), and (ii) during reappraisal in the beta band at left dlPFC, attachment anxiety and avoidance showed opposite associations (β = −0.13 and β = +0.12, respectively). Given the achieved sample size (N = 40), α = .05, and a repeated-measures ANOVA framework with three within-subject conditions, the design was sensitive to effects of f = 0.22 (η²p = .05), consistent with small-to-medium effects typically observed in EEG emotion studies (e.g. Uusberg et al. 2014).

Ethical approval was granted by the Scientific Ethics Committee of Universidad Católica del Norte (Resolutions No. 099/2021 and 037/2023). All procedures adhered to the Declaration of Helsinki and institutional ethical guidelines for human research.

### 2.2. Instruments

#### 2.2.1. Experiences in Close Relationships Questionnaire (ECR-12)

Attachment was evaluated using the Spanish adaptation of the 12-item Experiences in Close Relationships Questionnaire (ECR-12), validated for the Chilean population by Guzmán-González et al. (2020). This self-report measure assesses two dimensions of adult attachment: anxiety and avoidance, each represented by six items (e.g., “*I feel uncomfortable opening up to my partner*” for avoidance; “*If I cannot get my partner to show interest in me, I get upset or angry*” for anxiety) (Brennan et al., 1998). Participants responded on a 7-point Likert scale ranging from 1 (strongly disagree) to 7 (strongly agree), with higher scores reflecting greater attachment insecurity (Guzmán-González et al., 2023). In the present sample, the scale demonstrated satisfactory internal consistency (α = .76 for attachment anxiety, α = .84 for avoidance), aligning with previous psychometric findings for both the 12- and 36-item Chilean versions (Guzmán-González et al., 2020; Spencer et al., 2013).

### 2.3. Emotion Regulation Task

Emotion regulation was evaluated using an experimental paradigm adapted from Ochsner et al. (2004), Vrtička et al. (2012), and recent studies by Domic-Siede et al. (2024, 2025a, 2025b, 2025c). The task was implemented in *Presentation®* software (Version 18.0, Neurobehavioral Systems) and included 60 images from the International Affective Picture System (IAPS; Lang et al., 2005): 45 negative and 15 neutral, selected according to previous ER research (Moser et al., 2009; Schlumpf et al., 2019; Domic-Siede et al., 2023a) (see **Supplementary Tables S1–S3**).

Participants performed three experimental conditions—Natural, Reappraise, and Suppress (Domic-Siede et al., 2023a)—while stimuli were displayed on a 23.6″ ASUS VG248QE monitor. Prior to the main task, a training phase was conducted consisting of three blocks of three practice trials per condition, which included visual examples and instructions for using the Self-Assessment Manikin (SAM; Bradley & Lang, 1994) to rate emotional arousal.

In the Natural condition, participants simply viewed each image and allowed emotions to occur spontaneously. In Reappraise, they employed cognitive reappraisal strategies to reinterpret the image (e.g., imagining a positive or fictional outcome; Ochsner et al., 2004). In Suppress, they were instructed to inhibit any facial or bodily expression of emotion. Following each image, participants rated the intensity of their emotional experience on a 7-point SAM scale (1 = low, 7 = high) in response to the prompt: *“Indicate the intensity of your emotional response to the image you just saw.”* The task comprised 12 randomized blocks of five trials each. The Natural condition included 30 trials (15 neutral and 15 negative: Nat-neutral and Nat-negative), while Reappraise and Suppress each included 15 trials using only negative images. Each participant viewed the same fixed image set, with every image assigned exclusively to one condition (Natural, Reappraise, or Suppress). Both the block order and the trial sequence within blocks were randomized individually for each participant. Short breaks were provided between blocks to reduce fatigue.

Each trial followed a fixed temporal sequence (**Fig. 1**): (i) a 3-second fixation cross, (ii) a 2-second instruction cue, (iii) a 1-second fixation, (iv) a 4-second image presentation, and (v) the arousal rating screen. This sequence was repeated across all trials in all conditions.

**Figure 1.**
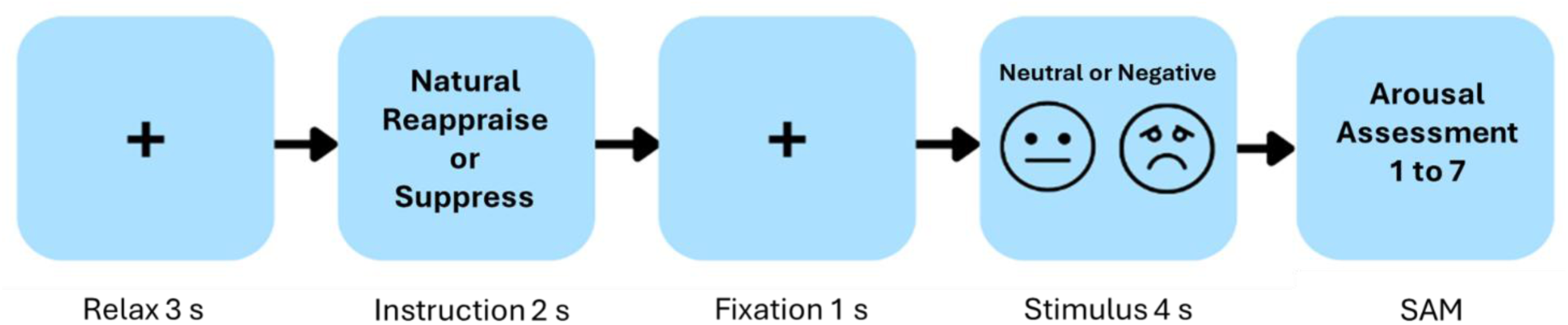
The diagram depicts the sequence of events within a representative trial of the experimental task used to examine how different emotion regulation strategies influence affective responses. Each trial starts with a 3-second relaxation period indicated by a fixation cross, followed by a 2-second cue instructing participants to apply one of three strategies: respond naturally, suppress their emotional expression, or reappraise the meaning of the upcoming stimulus. After a 1-second fixation interval, a neutral or negative image is presented for 4 seconds. Immediately afterward, participants report their perceived arousal on a 7-point Likert scale, where 1 denotes low emotional intensity and 7 denotes high emotional intensity. SAM = Self-Assessment Manikin.

**Figure 2.**
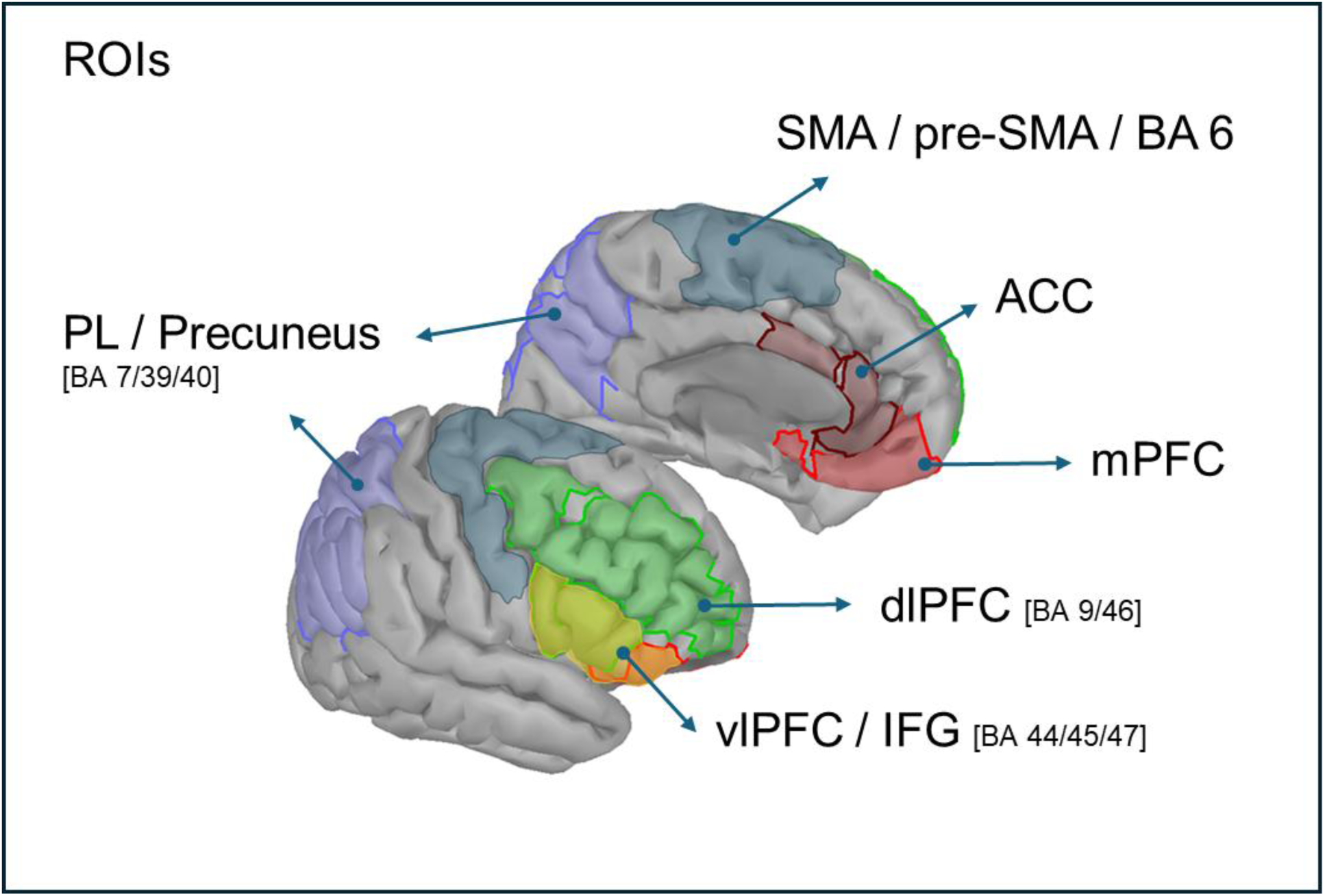
Regions of Interest (ROIs) selected for source-level analysis. Cortical regions of interest were defined according to the Destrieux atlas and bilateral Brodmann areas. ROIs included the anterior cingulate cortex (ACC), medial prefrontal cortex (mPFC), dorsolateral prefrontal cortex (dlPFC; BA 9/46), ventrolateral prefrontal cortex / inferior frontal gyrus (vlPFC/IFG; BA 44/45/47), supplementary and pre-supplementary motor areas (SMA/pre-SMA; BA 6), and parietal lateral cortex / precuneus (PL/Precuneus; BA 7/39/40). These regions were used to extract principal component time series (PCA) from the reconstructed source activity within the theta (4–8 Hz) and beta (15–30 Hz) frequency bands.

### 2.4. Electroencephalographic Data Acquisition

Electroencephalographic (EEG) activity was recorded using a BioSemiⓇ system (www.biosemi.com) equipped with 64 scalp electrodes and two mastoid electrodes, arranged according to the international 10/20 system (Keil et al., 2014). The signal was continuously sampled at 2048 Hz and referenced online to the Common Mode Sense (CMS) and Driven Right Leg (DRL) active electrodes. All electrode impedances were maintained below 20 kΩ throughout the recording.

### 2.6. Data Analysis

#### 2.6.1. Behavioral Data Analysis

Behavioural data were processed using *GraphPad Prism 8* and *MATLAB R2022b*. Descriptive statistics were computed to summarize participants’ self-reported arousal ratings. Data normality was evaluated with the Shapiro–Wilk test (Mishra et al., 2019), which indicated significant deviations and skewness; therefore, non-parametric analyses were employed.

To verify that the experimental manipulation produced the expected changes in emotional arousal, a Friedman test was conducted as the non-parametric equivalent of a repeated-measures ANOVA (Field, 2013). When significant main effects were observed, Dunn’s post-hoc comparisons were applied, and Cohen’s *d* was computed to estimate effect sizes for each pairwise contrast.

This analysis served as a manipulation check to confirm the overall task effect on arousal across conditions.

#### 2.6.2. EEG Signal Preprocessing

EEG data were preprocessed following established procedures (Domic-Siede et al., 2021, 2023b, 2024, 2025c) using EEGLAB toolbox (Delorme & Makeig, 2004). First, a high-pass finite impulse response (FIR) filter at 0.1 Hz was applied to remove slow drifts, followed by a low-pass filter at 40 Hz to attenuate high-frequency noise. The continuous data were then downsampled to 512 Hz.

Epochs were extracted from −1000 ms to 4000 ms relative to the onset of the IAPS image for four conditions: Natural Neutral, Natural Negative, Reappraise, and Suppress. Trials containing gross artifacts were visually excluded. Artifact correction was further refined through Independent Component Analysis (ICA) using the Logistic Infomax algorithm (Bell & Sejnowski, 1994). Components corresponding to non-neural sources (e.g., eye blinks, muscle activity, cardiac artifacts) were identified with ICLabel (Pion-Tonachini et al., 2019) and verified by visual inspection. This preprocessing pipeline maximized the signal-to-noise ratio while maintaining spatial accuracy for subsequent analyses.

#### 2.6.3. Brain Sources Reconstruction

Cortical source estimation was performed using the open-access software Brainstorm (Tadel et al., 2011), available under the GNU General Public License (http://neuroimage.usc.edu/brainstorm). Source reconstruction was applied to preprocessed EEG data (filtered between 0.1 and 40 Hz and artifact-free) corresponding to the 4-s image-viewing period in each trial. The sources were estimated using the Standardized Low-Resolution Brain Electromagnetic Tomography (sLORETA) algorithm (Pascual-Marqui, 2002), based on the ICBM152 anatomical template and the standard 10–20 electrode system. The minimum-norm imaging approach was implemented using the symmetric Boundary Element Method (BEM) forward model provided by OpenMEEG (Gramfort et al., 2010).

##### 2.6.3.1. Theta Band Source (4 –8 Hz)

Oscillatory power in the theta band was estimated directly from the reconstructed cortical sources. For each participant and condition, the power spectral density (PSD) was computed using the Welch method (1-s Hamming windows with 50% overlap). The PSD was calculated separately for the task period (0.3–4 s post-stimulus) and for the baseline period (−1 to 0 s). The 0.3–4 s window was selected *a priori* based on prior EEG studies of emotion regulation showing that regulatory engagement and sustained affective processing occur within this interval (e.g., Domic-Siede et al., 2024, 2025a, 2025b; Ertl et al., 2013; Hajcak & Nieuwenhuis, 2006; Moser et al., 2006). Theta power during the task was then normalized using a symmetric contrast between the task and baseline periods, defined as:

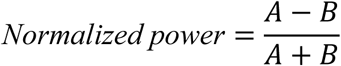

where *A* corresponds to the PSD during the task period and *B* to the PSD during the baseline period.

Regions of interest (ROIs) were anatomically defined using a multi-atlas convergence approach based on the Destrieux atlas (Destrieux et al., 2010), Brodmann areas (Fischl, 2013), the Desikan–Killiany–Tourville (DKT) atlas (Klein & Tourville, 2012), and the Desikan–Killiany atlas (Desikan et al., 2006). The following bilateral regions were selected: anterior cingulate cortex (ACC), dorsolateral prefrontal cortex (dlPFC), medial prefrontal cortex (mPFC), and ventrolateral prefrontal cortex / inferior frontal gyrus (vlPFC/IFG). For each ROI, a representative time series was extracted using Principal Component Analysis (PCA), and the first principal component was selected as the index of dominant theta activity (**Figure 2**).

**Figure 2a.**
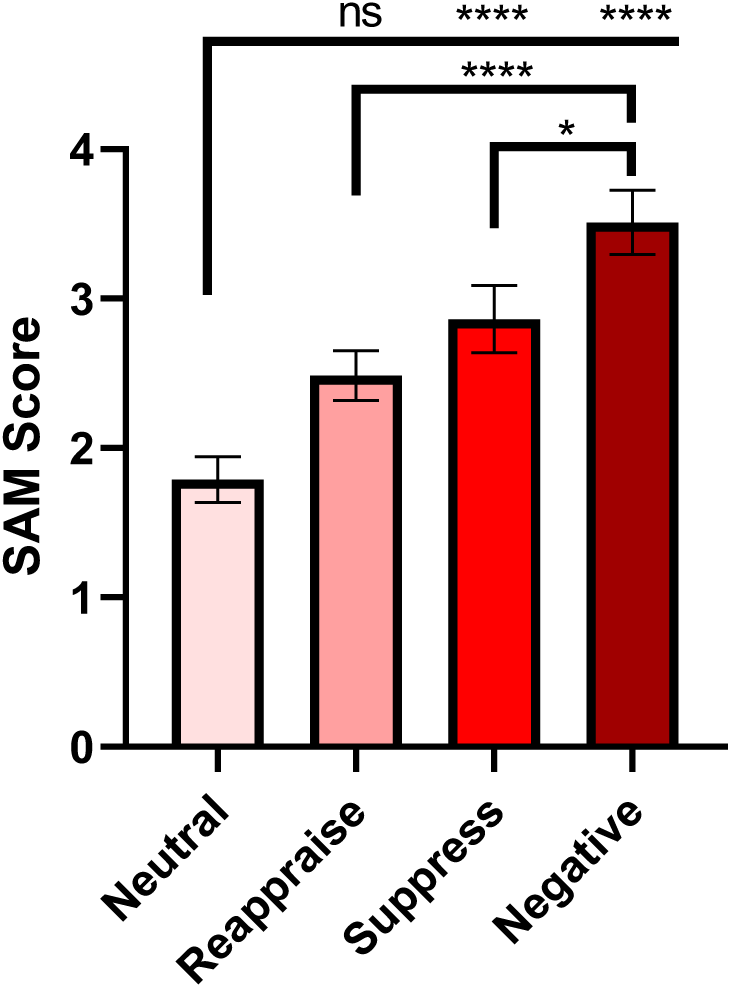
Self-reported arousal across emotion regulation conditions. Mean Self-Assessment Manikin (SAM) arousal ratings (1–7 scale) are shown for the four experimental conditions: Neutral, Reappraise, Suppress, and Negative. A Friedman test revealed a significant main effect of Condition, χ²(3) = 57.34, *p* < .0001. Post-hoc Dunn’s tests indicated higher arousal in Suppress (*p* < .0001, *d* = 0.97) and Negative (*p* < .0001, *d* = 1.57) relative to Neutral. Negative images also elicited greater arousal than Reappraise (*p* < .0001, *d* = 1.00) and Suppress (*p* = .038, *d* = 0.60). Error bars represent ±1 SEM

##### 2.6.3.2. Beta Band Source (15 –30 Hz)

The analysis of beta oscillatory activity followed the same procedure used for the theta band. PSD was computed with the Welch method in the 15–30 Hz range for both the task (0.3–4 s) and baseline (−1 to 0 s) periods. This temporal window was also defined a priori, consistent with previous work on emotion regulation (Domic-Siede et al., 2024, 2025a, 2025b; Ertl et al., 2013; Hajcak & Nieuwenhuis, 2006; Moser et al., 2006). Normalized beta power was obtained using the same symmetric contrast (A−B)/(A+B), with *A* corresponding to task PSD and *B* to baseline PSD.

The bilateral ROIs defined for beta analysis (derived from the same multi-atlas framework: Destrieux, Brodmann, DKT, and Desikan atlases) included the supplementary motor and premotor area (BA6), parietal lateral cortex / precuneus (PL/Precuneus), dorsolateral prefrontal cortex (dlPFC), and ventrolateral prefrontal cortex / inferior frontal gyrus (vlPFC/IFG). As in the theta analysis, PCA was applied within each ROI, and the first component was used as the representative measure of dominant beta activity. Normalized PSD maps were averaged across participants for each condition. Group-level cortical maps were spatially normalized and projected onto the ICBM152 template surface for visualization. Cortical activations were displayed using Brainstorm’s default colormap and threshold settings, illustrating the spatial distribution of task-related power changes in the theta and beta frequency bands.

#### 2.6.3. Linear Mixed-Effects Models

To test the hypotheses regarding the modulation of oscillatory power by **Condition** (Negative, Reappraise, Suppress), **Region of Interest (ROI)**, and affective covariates (**Attachment Anxiety [ANX]** and **Attachment Avoidance [AVD]**), we fitted **linear mixed-effects models (LMMs)** separately for the **theta** and **beta** frequency bands using MATLAB (`fitlme`, maximum likelihood estimation).

##### Data preprocessing and coding

- **Subject** was treated as a random factor.
- **Condition** had three levels (`Negative`, `Reappraise`, `Suppress`), with **Negative** as the reference.
- **ROI** was modeled as a categorical fixed factor, with different sets for each frequency band:

`o` **Theta band:** L/R ACC, L/R dlPFC, L/R mPFC, L/R vlPFC/IFG (reference = *Left vlPFC/IFG*).
`o` **Beta band:** L/R BA6, L/R PL/Precuneus, L/R dlPFC, L/R vlPFC/IFG (reference = *Left BA6*).
- Continuous covariates (**ANX** and **AVD**) were **mean-centered** at the sample level: `ANX_c`, `AVD_c`.
- The dependent variable was **Amplitude** (log-transformed, normalized oscillatory power).

##### Fixed-effects structure

Each model included all main effects and their interactions up to three-way terms among Condition, ROI, and the covariates:

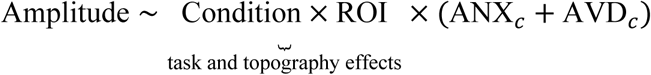

This expands to:

- Main effects: Condition, ROI, ANX_c, AVD_c
- Two-way interactions: Condition×ROI, Condition×ANX_c, ROI×ANX_c, Condition×AVD_c, ROI×AVD_c
- Three-way interactions: Condition×ROI×ANX_c and Condition×ROI×AVD_c

This structure allowed us to test whether **Reappraise** increased theta activity in frontal ROIs, and whether **higher attachment anxiety** attenuated this effect (Condition×ROI×ANX_c), as well as analogous modulations by **AVD**.

##### Random-effects structure

To account for inter-individual variability, we specified random intercepts and slopes for Condition by subject, as well as ROI-specific intercepts nested within each subject:

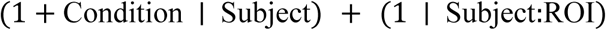

This structure models (a) subject-level variation in both intercepts and condition slopes (and their correlations), and (b) ROI-level variability within subjects.

##### Inference and reporting

- **Degrees of freedom** and **p-values** were estimated using the **Satterthwaite approximation** (default in `fitlme`).
- Fixed effects were reported as β (SE), t, p, and 95% confidence intervals.
- **Model fit indices** included AIC, BIC, and log-likelihood values.
- **Simple slopes** and **planned contrasts** (e.g., effect of `ANX_c` within *Reappraise × R dlPFC*) were tested using `coefTest`, with marginal predictions (`predict`) at ±1 SD of ANX_c or AVD_c.

##### Robustness and influence diagnostics

Residuals, leverage values, and **Cook’s distance** were examined for influential subjects or ROI-level observations. We also verified the stability of key effects (e.g., *Condition×ROI×ANX_c* in the right dlPFC, theta band) through **leave-one-subject-out** analyses. Main results are reported along with these checks when they affected inference.

#### 2.6.4. Spearman Rank Correlations

To complement the mixed-effects models, we conducted non-parametric Spearman rank correlations to examine bivariate associations among behavioral and neural variables. These analyses were designed to assess the functional significance of the cortical sources identified in the LMMs as showing the strongest task- and trait-related effects. Specifically, only the right dlPFC (theta band) and left dlPFC (beta band) were selected, as these regions and frequency bands exhibited significant or theoretically meaningful modulations by condition and attachment orientations in the multilevel analyses (see Section 3.3).

Beyond their neural interpretation, these correlations also served to explore whether individual differences in attachment orientations were reflected in subjective emotional experience. This approach provided a complementary behavioral index testing whether anxious and avoidant attachments were associated with differential emotional reactivity and regulation success across conditions. This targeted analysis therefore allowed us to determine (a) whether the magnitude of source activity covaried with subjective arousal, and (b) whether attachment-related traits mapped onto individual differences in emotional intensity across emotion regulation contexts.

##### Neural–behavioral correlations

1. Right dlPFC theta (4–8 Hz) × arousal: Following the a priori effect tested in the LMMs, we correlated normalized theta power from the right dlPFC ROI with self-reported arousal separately for each condition (Neutral, Reappraise, Suppress, Negative).
2. Left dlPFC beta (15–30 Hz) × arousal: Motivated by the left frontal beta effects in the LMMs, we correlated normalized beta power from the left dlPFC ROI with arousal separately for each condition.

##### Behavior–trait correlations

We also examined the association between self-reported arousal and attachment dimensions (ECR-12 Anxiety [ANX], Avoidance [AVD]) within each condition to test whether individual differences in attachment mapped onto perceived emotional intensity.

## 3. Results

### 3.1. Behavioral Results

Self-reported arousal ratings (1–7 scale) were compared across four conditions: Natural Neutral, Reappraise, Suppress, and Natural Negative. Descriptive analyses revealed a clear progressive increase in arousal from Natural Neutral (M = 1.79, SD = 0.97) to Natural Negative (M = 3.51, SD = 1.37), with intermediate values for Reappraise (M = 2.49, SD = 1.05) and Suppress (M = 2.86, SD = 1.42) (Supplementary **Table S4**). Although the Shapiro–Wilk test indicated that Reappraise data did not deviate from normality (*p* = .099), other conditions showed non-normal distributions (*ps* < .05; Supplementary **Table S5**). Therefore, a nonparametric Friedman test was applied, revealing a significant main effect of Condition, χ²(3) = 57.34, *p* < .0001 (Supplementary **Table S6**).

Post-hoc Dunn’s multiple comparisons (**Supplementary Tables S7–S8**) indicated significantly higher arousal in Suppress (*p* < .0001, *d* = 0.97) and Natural Negative (*p* < .0001, *d* = 1.57) compared to Natural Neutral. The Reappraise condition did not differ significantly from Neutral (*p* = .0638), nor from Suppress (*p* = .4138). However, Reappraise and Suppress both elicited lower arousal than Natural Negative (*p*s < .0001, *d*s = 1.00 and 0.60, respectively; **Figure 2**).

These results indicate that the Suppress and Natural Negative conditions produced markedly greater perceived arousal relative to the neutral baseline. In contrast, although Reappraise tended to increase arousal compared to Neutral, its levels remained lower than those in the Natural Negative condition, consistent with its regulatory effectiveness.

These analyses served as a manipulation check to confirm the expected condition effect on arousal.

### 3.2. Brain Sources Reconstruction Results

#### 3.2.1. Global Source Reconstruction in Theta Band (4–8 Hz) – *Reappraise* Condition

The cortical distribution of theta power (4–8 Hz) during the Reappraise condition showed clear positive contrasts (*A* − *B*)/(*A* + *B*)in frontal and midline regions compared with baseline (−1 to 0 s). The strongest theta activity appeared in the right dorsolateral prefrontal cortex (dlPFC), right ventrolateral prefrontal cortex/inferior frontal gyrus (vlPFC/IFG), and right medial prefrontal cortex (mPFC), with additional increases in posterior parietal cortex and precuneus. This spatial pattern indicates predominant fronto-posterior engagement consistent with the recruitment of top-down control networks typically linked to cognitive reappraisal. Figure 3 shows the normalized theta source maps projected on the ICBM152 template for the 0.3–4 s post-stimulus interval.

**Figure 3.**
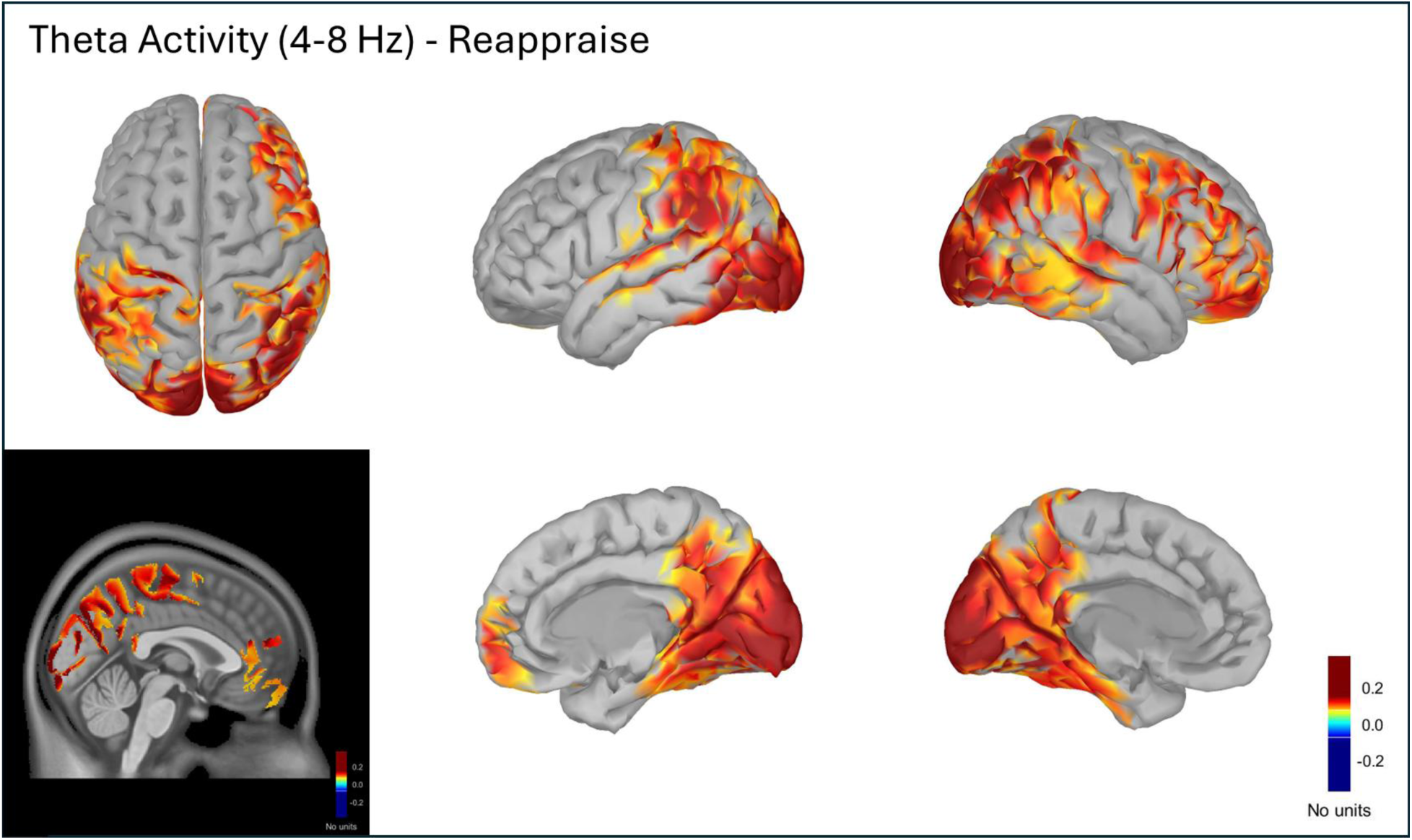
Cortical theta (4–8 Hz) source activity during the Reappraise condition. Normalized power spectral density (PSD) maps computed using the (*A* − *B*)/(*A* + *B*) contrast between task (A) and baseline (B) periods. PSD values for A and B were obtained with the Welch method in physical units of *μV*^2^/*Hz*; the normalization yielded unitless ratios representing relative changes in spectral power. The figure displays averaged theta activity from 0.3 to 4 s post-stimulus, projected onto the ICBM152 cortical surface. Warmer colors indicate increased theta power relative to baseline, predominantly in the right dorsolateral prefrontal cortex (dlPFC), ventrolateral prefrontal cortex/inferior frontal gyrus (vlPFC/IFG), medial prefrontal cortex (mPFC), and posterior parietal regions including the precuneus.

#### 3.2.2. Global Source Reconstruction in Beta Band (15–30 Hz) – *Suppress* Condition

The Suppress condition displayed a more heterogeneous modulation of beta power (15–30 Hz) relative to baseline, with both positive and negative contrasts across the cortex. Decreases in beta activity (blue-coded) were visible in portions of the left dlPFC (BA 9/46), right vlPFC/IFG (BA 44/45), bilateral SMA/pre-SMA, bilateral mPFC, and parts of the posterior parietal cortex. Conversely, increases (warm-coded) were observed over the bilateral precuneus (BA 7/39/40), posterior parietal and occipital cortices, bilateral mPFC, and right dlPFC.

This mixed configuration suggests differentiated fronto-parietal engagement: posterior and prefrontal beta enhancements possibly supporting higher-order control and self-monitoring, together with central beta decreases associated with reduced motor readiness during expressive suppression. **Figure 4** depicts these normalized beta maps across multiple cortical views within the 0.3–4 s window.

**Figure 4.**
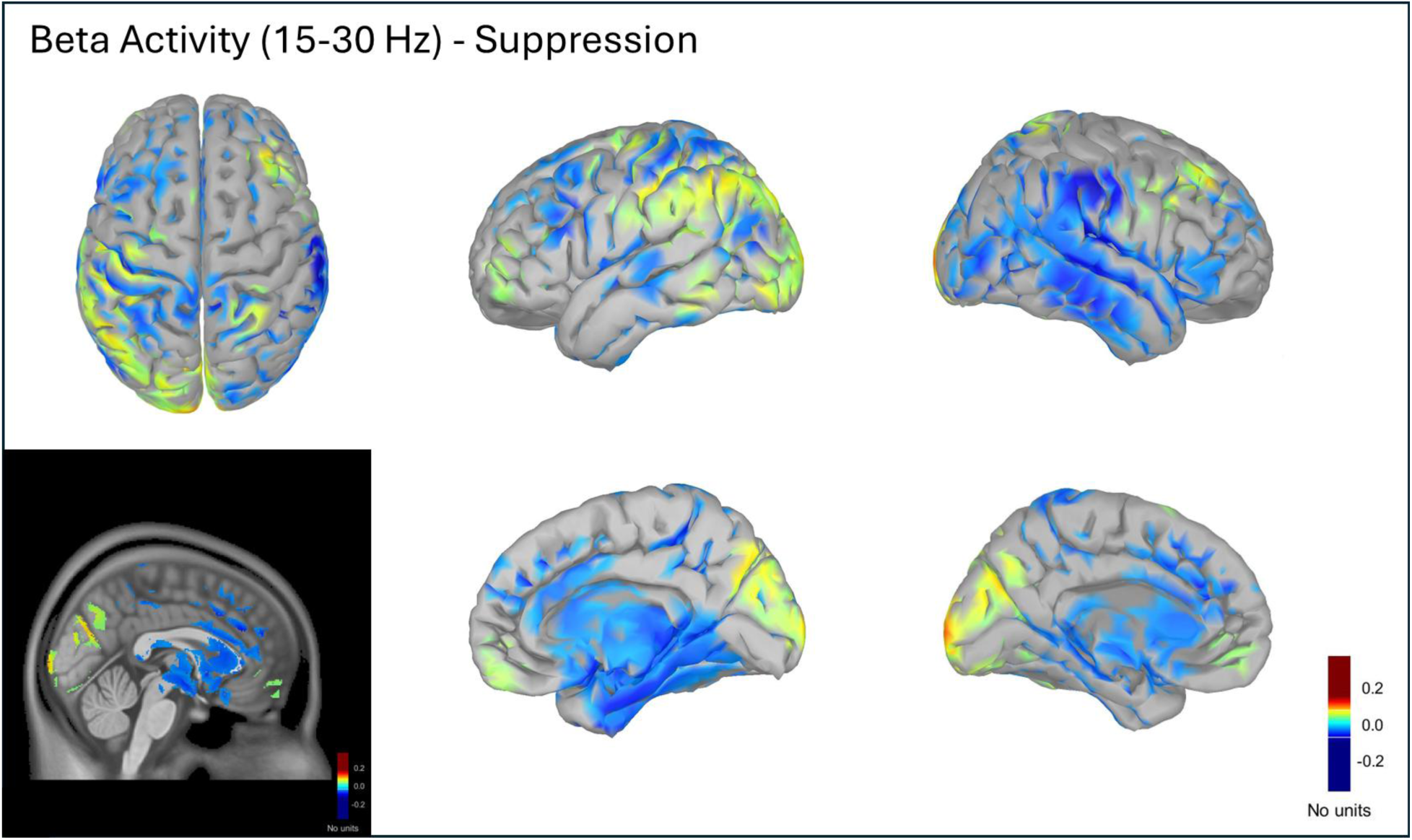
Cortical beta (15–30 Hz) source activity during the Suppress condition. Normalized power spectral density (PSD) maps computed using the (*A* − *B*)/(*A* + *B*) contrast between task (A) and baseline (B) periods. PSD values for A and B were estimated with the Welch method in physical units of *μV*^2^/*Hz*; the normalization produced unitless ratios reflecting relative changes in spectral power. The figure displays averaged beta activity from 0.3 to 4 s post-stimulus, projected onto the ICBM152 cortical surface. Cooler colors (negative contrasts) indicate decreased beta power relative to baseline, particularly across posterior parietal cortex, precuneus, and medial frontal regions, whereas warmer colors (positive contrasts) mark localized increases in posterior and dorsolateral prefrontal areas.

### 3.3. Linear Mixed-Effects Models Results

We fit separate LMMs for theta and beta power (see Methods for structure). Both models converged under ML and showed acceptable residual dispersion.

#### Model fit

The theta model (reference: Negative × L vlPFC/IFG) yielded AIC = 492.09, BIC = 881.44, log-likelihood = −166.04. The beta model (reference: Negative × L BA6) yielded AIC = −1.27, BIC = 388.09, log-likelihood = 80.64 (**Table 2**).

**Table 2.**
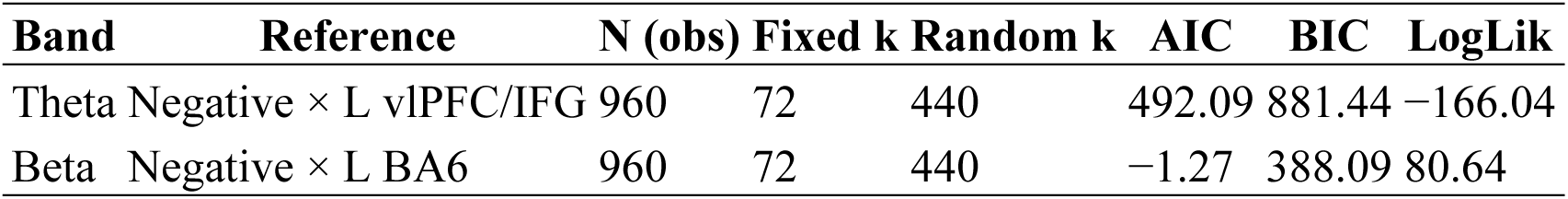
Model fit indices.

#### Theta band

There were no omnibus main effects of Condition across ROIs (ps >.23). Critically, we observed a Condition × ROI × ANX interaction specific to the right dlPFC during Reappraise: higher attachment anxiety predicted lower theta power (β = −0.187, SE = 0.075, t = −2.48, p = .013). This pattern is consistent with the hypothesis that anxiety attenuates frontal theta engagement during cognitive reappraisal (**Table 3**, Supplementary **Table S9** and **S10**).

**Table 3.** Theta LMM — effects relevant to hypotheses Effect (theta) β SE t p.

| Effect (theta) | $\beta$ | SE | t | p |
| --- | --- | --- | --- | --- |
| Condition: Reappraise | 0.024 | 0.066 | 0.37 | .714 |
| Condition: Suppress | 0.082 | 0.069 | 1.20 | .232 |
| ROI: R dlPFC | 0.107 | 0.059 | 1.81 | .071 |
| Reappraise × R dlPFC × ANX | **-0.187** | **0.075** | **-2.48** | **.013** |
| Suppress × L mPFC × AVD | 0.139 | 0.069 | 2.00 | **.046** |
| Suppress × R mPFC × AVD | 0.140 | 0.070 | 2.09 | **.045** |
| Reappraise × ROI × AVD (any frontal) | — | — | — | $\geq .19$ |
| Reappraise × ROI × ANX (others) | — | — | — | $\geq .15$ |

Attachment avoidance (AVD) did not significantly modulate theta during Reappraise (ps ≥ .19), but showed a *trend toward increased theta* during Suppress in the mPFC bilaterally (L mPFC: β = 0.139, SE = 0.069, t = 2.00, p = .046; R mPFC: β = 0.140, SE = 0.069, t = 2.01, p = .045). The right dlPFC also exhibited a trend toward higher theta overall (β = 0.107, SE = 0.059, p = .071) (**Table 2**, Supplementary **Table S9** and **S10**).

Random effects indicated substantial between-subject variability in condition slopes (Reappraise SD = 0.197; Suppress SD = 0.232) with negative intercept–slope correlations (see Supplementary **Table S11**).

#### Beta band

No omnibus Condition main effects emerged (ps ≥ .32). Task × ROI effects showed lower beta during Suppress in right PL/Precuneus (β = −0.216, SE = 0.067, t = −3.25, p = .001, Supplementary Figure S1), alongside a positive main effect for right PL/Precuneus (β = 0.113, SE = 0.048, p = .018) (Table 3, Supplementary **Table S12**).

Affective covariates modulated beta selectively in left dlPFC during Reappraise: ANX was associated with lower beta (β = −0.127, SE = 0.061, t = −2.08, p = .038), whereas AVD was associated with higher beta (β = 0.124, SE = 0.056, t = 2.21, p = .027). No other ANX/AVD interactions reached significance (ps ≥ .08) (**Table 4**, Supplementary **Table S13**).

**Table 4.** Beta LMM — effects relevant to hypotheses.

| **Effect (beta)** | **$\beta$** | **SE** | **t** | **p** |
| --- | --- | --- | --- | --- |
| Condition: Reappraise | 0.050 | 0.050 | 1.00 | .315 |
| Condition: Suppress | 0.034 | 0.048 | 0.70 | .483 |
| ROI: R PL/Precuneus | 0.113 | 0.048 | 2.38 | .018 |
| Suppress × R PL/Precuneus | **-0.216** | **0.067** | **-3.25** | **.001** |
| Reappraise × L dlPFC × ANX | **-0.127** | **0.061** | **-2.08** | **.038** |
| Reappraise × L dlPFC × AVD | **0.124** | **0.056** | **2.21** | **.027** |

In summary, reappraisal-related frontal theta was attenuated by higher attachment anxiety in the right dlPFC, whereas attachment avoidance showed only a trend-level increase in theta during suppression in the mPFC. For beta power, suppression decreased beta in the right parietal lobe and precuneus, while during reappraisal, attachment anxiety decreased and attachment avoidance increased beta in the left dlPFC.

### 3.4. Correlations Between dlPFC Theta Power and Self-Reported Arousal

To further examine the functional relevance of the right dorsolateral prefrontal cortex (R dlPFC) theta activity observed during the Reappraise condition (**Figure 5A**), we tested whether self-reported arousal levels in each condition were associated with theta power extracted from this ROI. In addition, these analyses provided an opportunity to examine whether subjective emotional responses varied as a function of attachment dimensions, serving as a behavioral correlate of the neural effects and offering indirect evidence that individuals with different attachment orientations experience and regulate emotions differently.

**Figure 5.**
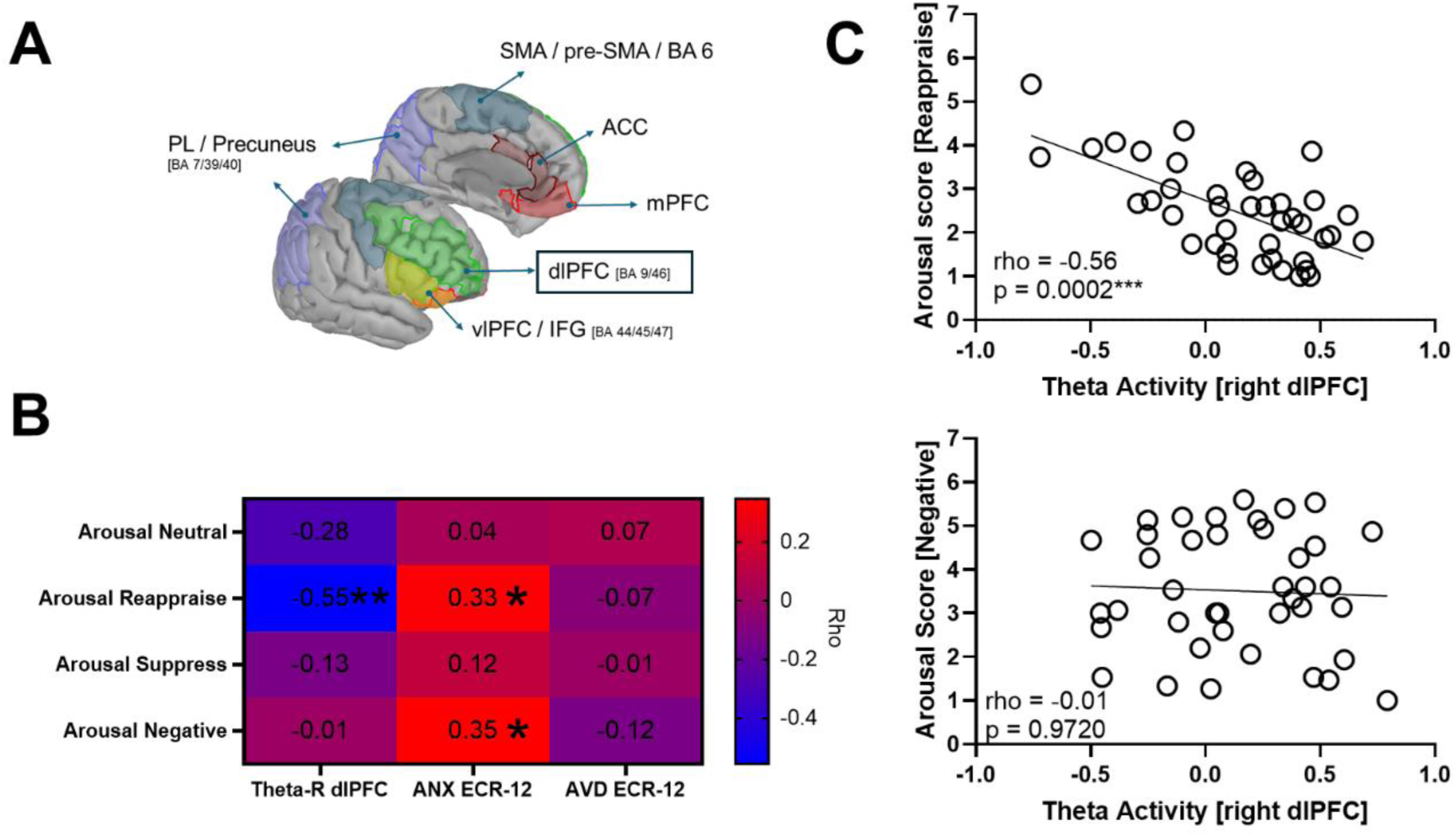
Associations between right dlPFC theta activity, self-reported arousal, and attachment orientations. (A) Cortical localization of the right dorsolateral prefrontal cortex (R dlPFC) region of interest (ROI) used for correlation analyses, displayed on the ICBM152 cortical template. (B) Heatmap showing Spearman’s rho coefficients between theta power in the R dlPFC (first column), attachment anxiety (ANX; second column), and attachment avoidance (AVD; third column) with self-reported arousal across the four experimental conditions (rows). Blue hues denote negative correlations and red hues positive correlations; *p* < .05 (*), \**p* < .01 (**). (C) Scatter plots illustrating the negative association between R dlPFC theta power and arousal during the Reappraise condition (top) and the absence of correlation during the Negative condition (bottom).

Spearman correlations revealed a significant negative association between theta power in the R dlPFC and self-reported arousal during the Reappraise condition (rho = −0.55, p = 0.0002), indicating that greater frontal theta activity was linked to lower perceived arousal while reappraising negative stimuli (**Figure 5B**, **Figure 5C**, top). No significant correlations emerged between theta and arousal in the Neutral (rho = −0.28, p = 0.076), Suppress (rho = −0.13, p = 0.435), or Negative (rho = −0.01, p = 0.972) conditions (**Figure 5B**, **Figure 5C**, bottom).

Regarding attachment dimensions, attachment anxiety correlated positively with arousal during both Reappraise (rho = 0.33, p = 0.036) and Negative (rho = 0.35, p = 0.028) conditions, suggesting that individuals with higher attachment anxiety reported greater emotional activation in response to negative stimuli and during attempts to regulate them. In contrast, no significant associations were found between arousal and attachment avoidance across any condition (all ps > .44).

### 3.5. Correlations Between Left dlPFC Beta Power and Self-Reported Arousal

To examine the functional significance of left dorsolateral prefrontal cortex (L dlPFC) beta modulation observed during the Reappraise condition, we correlated mean beta power (15–30 Hz) extracted from this ROI with self-reported arousal across conditions. Spearman tests (N = 40) showed a **specific negative association under Natural Negative**: higher L dlPFC beta related to **lower** arousal (rho = −0.340, 95% CI [−0.595, −0.022], *p* = .032). No other condition yielded reliable effects (Reappraise: rho = −0.011, 95% CI [−0.330, 0.310], *p* = .947; Neutral: rho = 0.151, 95% CI [−0.178, 0.450], *p* = .352).

## 3. Discussion

The present findings reveal that adult attachment orientations are reflected in distinct patterns of cortical oscillatory dynamics during emotion regulation. Although cognitive reappraisal did not elicit a significant overall theta power increase, individuals with higher attachment anxiety exhibited reduced theta-band activity in the right dorsolateral prefrontal cortex (dlPFC) during reappraisal, suggesting attenuated recruitment of cognitive control mechanisms (Domic-Siede et al., 2024; Ertl et al., 2013; Zhang et al., 2025; Zhao et al., 2021). In contrast, expressive suppression was characterized by decreased beta power in the right parietal cortex and precuneus—regions involved in self-referential attention—consistent with attentional disengagement from self-related processing (Northoff et al., 2006; Ochsner et al., 2012; Spitzer & Haegens, 2017).

During reappraisal, attachment anxiety was linked to lower beta power and attachment avoidance to higher beta power in the left dlPFC. These results indicate that reappraisal and suppression engage distinct oscillatory systems and that attachment orientations modulate their neural implementation. Frontal theta activity indexes top-down control processes during reappraisal, whereas beta oscillations reveal complementary functions: frontal beta reflects regulatory control differences associated with attachment (Abid et al., 2024; Domic-Siede et al., 2025b; Etkin et al., 2015; Feeser et al., 2014; He et al., 2023; Long et al., 2020; Zhao et al., 2021), and parietal beta decreases during suppression suggest reduced self-referential attention (Northoff et al., 2006; Ochsner et al., 2012).

Frontal midline theta is a well-established marker of cognitive control engagement, particularly during emotion regulation (Cavanagh & Frank, 2014), while beta-band oscillations in fronto-parietal networks sustain ongoing cognitive or sensorimotor states by inhibiting change (Engel & Fries, 2010; Spitzer & Haegens, 2017). From this broader perspective, reduced frontal theta in higher attachment anxiety individuals may reflect difficulty in recruiting executive control under emotional stress—consistent with heightened reactivity and low regulatory flexibility (Diamond & Fagundes, 2010; Mikulincer & Shaver, 2019; Silva, 2005). Conversely, enhanced frontal beta in avoidant individuals could indicate rigid top-down control supporting emotional detachment (Knyazev, 2007; Mikulincer & Shaver, 2019). Thus, insecure attachment may map onto extremes of regulatory control: under-engagement in attachment anxiety and over-engagement in attachment avoidance.

Adopting an attachment framework, our oscillatory results provide neural evidence for these distinct regulatory tendencies. Higher attachment anxiety individuals showed reduced right-dlPFC theta during reappraisal, reflecting inefficient recruitment of cognitive control and a “hyperactivating” regulatory style easily overwhelmed by limbic-driven arousal (Domic-Siede et al., 2024; Long et al., 2020; Gillath et al., 2005; Mikulincer & Shaver, 2019; Vrtička & Vuilleumier, 2012). Higher attachment avoidance individuals, in turn, exhibited higher left-dlPFC beta, consistent with a “deactivating” style emphasizing cognitive distancing and suppression (Dörfel et al., 2014; Vrtička & Vuilleumier, 2012; Engel & Fries, 2010; Stoll et al., 2016). Together, these findings reveal opposite deviations in the recruitment of frontal control-related oscillations: Higher attachment anxiety individuals under-engaged theta-mediated control, while higher attachment avoidance individuals over-engaged beta-mediated inhibition. These complementary signatures for “hyperactivating” versus “deactivating” tendencies clarify how attachment orientations correspond to distinct neural mechanisms of regulation (Domic-Siede et al., 2025a, 2025b; Mikulincer & Shaver, 2019).

Bowlby’s concept of internal working models (IWMs) posits that early relationships shape emotion regulation strategies (Bowlby, 1969, 1973, 1982; Cassidy & Shaver, 2016). Our findings provide neurophysiological support: secure versus insecure orientations produced distinct neural responses during regulation. An anxious IWM—marked by fear of rejection—coincided with weak engagement of frontal control, whereas an avoidant IWM—marked by emotional deactivation—was associated with stronger prefrontal control. This aligns with evidence that attachment orientations are biologically embedded in emotion-related brain systems (Vrtička & Vuilleumier, 2012; Long et al., 2020; White et al., 2023).

Consistent with the Social Neuroscience of Attachment (SoNeAt) perspective (Vrtička et al., 2017; White et al., 2023), attachment orientations corresponded to differential recruitment of oscillatory networks. Behaviorally, higher attachment anxiety individuals tend to hyperactivate emotions, while avoidant individuals rely on suppression and withdrawal (Shaver & Mikulincer, 2007; Mikulincer & Shaver, 2019). Our electrophysiological findings mirror these profiles: lower frontal theta (under-regulation) in higher attachment anxiety and higher frontal beta (over-regulation) in higher attachment avoidance (Mikulincer & Shaver, 2019; Vrtička & Vuilleumier, 2012). Thus, attachment insecurity biases both emotional appraisal and regulation implementation, reflected here in frontal theta under-engagement and frontal beta over-engagement. Integrating biological levels of analysis into attachment theory (Thompson et al., 2022), our findings suggest that early relational experiences leave measurable neurophysiological imprints. Higher attachment anxiety individuals—with internal models of inconsistent support—showed vigilance and reduced top-down control, whereas higher attachment avoidance individuals—with models of self-reliance—showed enhanced inhibitory control and emotional disengagement. These results illustrate how IWMs are instantiated in the brain’s regulatory circuitry.

Moreover, our results align with the Neuro-Anatomical Model of Attachment (NAMA; Long et al., 2020), which proposes that attachment anxiety involves a hyper-reactive limbic system and reduced prefrontal regulation, whereas avoidance entails attenuated emotional appraisal and increased control. The observed reduction in right-dlPFC theta among anxious individuals and elevation of left-dlPFC beta among avoidant individuals fit this model (Klimesch, 2012; Vrtička & Vuilleumier, 2012; Cassidy & Shaver, 2016; Mikulincer & Shaver, 2016), offering empirical support for NAMA’s framework.

Finally, by capturing oscillatory markers of attachment-related regulation, this study bridges behavioral and neurophysiological evidence. Recent ERP studies show that attachment anxiety is linked to larger late positive potential (LPP) amplitudes during emotional processing (Domic-Siede et al., 2025a). Our source-level frequency findings complement these results, suggesting that weaker frontal theta engagement may underlie prolonged emotional reactivity in higher-anxiety individuals. Moreover, recent connectivity analyses have demonstrated that attachment orientations also modulate large-scale synchronization during emotion regulation, with anxious attachment showing reduced fronto-parietal theta coupling and avoidant attachment showing enhanced beta-band connectivity (Domic-Siede et al., 2025b). Together, these results indicate that IWMs are reflected not only in local oscillatory dynamics but also in the coordination of distributed regulatory networks, linking interpersonal dispositions to neural processes and advancing an integrative understanding of attachment and emotion regulation.

### 4.1. Clinical Implications

Difficulties in emotion regulation are transdiagnostic features present in a wide range of clinical conditions, including anxiety, depression, personality disorders, and trauma-related syndromes (Aldao et al., 2010; Cludius et al., 2020). Understanding the neurophysiological mechanisms that underpin regulatory strategies can therefore inform more precise and individualized interventions. The present findings highlight that individuals with different attachment orientations may engage distinct regulatory modes at the neural level, suggesting the potential clinical value of tailoring interventions accordingly.

Frontal theta and prefrontal beta activity appear to index different modes of emotion regulation, converging with long-standing clinical observations that anxiously attached individuals often experience emotional overwhelm, whereas avoidantly attached individuals rely more on detachment or suppression (Shaver & Mikulincer, 2002; Mikulincer & Shaver, 2019). At the neural level, diminished right dlPFC theta during reappraisal suggests that anxious individuals under-engage prefrontal control, which may explain why cognitive strategies like reframing/reappraisal often feel ineffective. In contrast, increased left dlPFC beta in avoidant individuals indicates sustained cognitive control or distancing, consistent with a rigid regulatory style aimed at minimizing affective engagement.

These patterns imply differentiated therapeutic approaches. For anxious clients, interventions that strengthen prefrontal regulatory engagement—such as cognitive reappraisal training, mindfulness, or neurofeedback targeting frontal theta—could enhance their capacity for top-down control. Preliminary evidence shows that neurofeedback protocols aimed at increasing frontal theta improve emotional regulation and reduce stress-related reactivity (Kang et al., 2014; Ros et al., 2013). For avoidant clients, therapy may focus on reducing excessive control and fostering safe emotional expression. Psychotherapeutic approaches such Emotionally focused Individual Therapy (Brubacher, 2017) or attachment-based therapies can help them notice their tendency to withdraw and gradually tolerate emotional activation (Fonagy & Bateman, 2006). By reframing high beta activity as a neural marker of overcontrol, therapists can use psychoeducation or biofeedback to promote awareness and relaxation during emotionally charged moments.

These findings highlight that attachment-related regulation patterns have identifiable neural correlates, suggesting that the habitual emotional responses of anxiously and avoidantly attached individuals are instantiate d in distinct brain dynamics that can nonetheless be modified through targeted intervention and relational experience. Communicating this to clients can reduce self-blame and enhance engagement: anxious individuals may feel validated knowing their difficulty in calming down has a neural basis that can change with practice, while avoidant individuals may begin to view their detachment as a learned neural habit that can be softened through relational safety.

More broadly, integrating these neural insights into attachment-based interventions could enrich psychoeducation in individual, couple, or group therapy. Understanding that anxious “hyperactivation” and avoidant “deactivation” correspond to distinct neural patterns may help clients and partners move from criticism to self-compassion—recognizing that these reactions are brain-based strategies for emotional protection that can be modified through therapeutic experience.

### 4.2. Limitations and Future Directions

Several limitations should be noted when interpreting these findings. Methodologically, the exclusive reliance on EEG constrains spatial precision. Although source reconstruction improves anatomical inference, deep or subcortical structures involved in emotion regulation—such as the amygdala, hippocampus, and limbic regions overall—cannot be accurately localized (Baillet, 2017). Multimodal approaches combining EEG and fMRI could clarify whether frontal theta activity during reappraisal co-occurs with BOLD activation in prefrontal control regions and reduced limbic reactivity, as observed in prior regulation research (Knyazev, 2012; Wager et al., 2008). Simultaneous EEG–fMRI could therefore strengthen the spatial validity of oscillatory interpretations.

Second, the study’s cross-sectional and correlational design limits causal inference. Attachment orientations and neural activity were measured at a single time point; thus, we cannot determine whether attachment orientations shape neural oscillations or vice versa. Longitudinal and experimental studies—such as attachment security priming (Gillath & Karantzas, 2019)—could examine whether temporary or enduring shifts in attachment security alter theta and beta dynamics during emotion regulation. Demonstrating such causal modulation would substantiate the functional role of attachment in shaping neural control processes.

Third, attachment was assessed using the self-report *Experiences in Close Relationships* (ECR) scale (Brennan et al., 1998), which captures conscious attachment tendencies but not deeper representational aspects. Interview-based measures like the *Adult Attachment Interview* (AAI; Main et al., 2008) or narrative tasks may index defensive processes more directly related to emotion regulation (Vrtička & Vuilleumier, 2012). Future studies integrating multiple assessment levels—self-report, representational, and behavioral—would help clarify which components of attachment predict neural variability. For instance, do AAI “dismissing” individuals exhibit the same beta enhancement as self-reported avoidants? Such convergence would strengthen construct validity and rule out method variance.

Conceptually, the paradigm involved regulating emotions elicited by unpleasant but impersonal IAPS images. Although this enhances experimental control, the absence of interpersonal cues means the attachment system may not have been fully engaged; attachment processes are most strongly activated in relational contexts (Mikulincer & Shaver, 2019). Thus, eliciting emotions without interpersonal content could potentially misrepresent or attenuate attachment-related effects. Investigating oscillatory activity during socially or attachment-relevant tasks—such as regulating emotions in response to partner feedback, rejection, or separation cues (Gillath et al., 2005)—would improve ecological validity. Combining neural data with autonomic indices could also reveal whether insecure attachment involves parallel physiological dysregulation (Diamond & Fagundes, 2010).

Cultural and developmental generalizability likewise warrant attention. The current mostly Chilean adult sample limits cross-cultural inference. Emotion regulation and attachment expression vary across societies (Mesquita et al., 2016); in cultures where restraint is normative, even secure individuals might show stronger beta activity during suppression. Comparative studies across individualistic and collectivistic contexts could clarify the cultural embedding of attachment-related neural patterns. Developmentally, longitudinal work tracking oscillatory activity from adolescence into adulthood could reveal when attachment-linked neural differences crystallize (Silvers et al., 2015; McRae et al., 2012). Such work would inform interventions aimed at promoting regulatory flexibility during sensitive periods of brain maturation.

Finally, future research should integrate behavioral, physiological, and computational levels of analysis (Petter et al., 2025). Emotion regulation is multifaceted, involving subjective, expressive, and neural components (Gross, 2015). Linking oscillatory indices with behavioral success (e.g., reductions in subjective arousal or facial expression) would test whether frontal theta predicts regulatory efficacy or beta reflects suppression effort even during reappraisal. Building on work that decodes affect from brain activity and demonstrates cross-modal correspondences (e.g., multivoxel “neural signatures” and ERP–fMRI links; Chang et al., 2015; Bo et al., 2021; Liu et al., 2012; Sabatinelli et al., 2013), and on studies where connectivity and task-evoked signals track regulation success and therapy-related change (Morawetz et al., 2017; Goldin et al., 2013; Dixon et al., 2020), multimodal, machine-learning pipelines are well-positioned to yield composite attachment-regulation profiles with predictive value for vulnerability and treatment responsiveness.

In summary, addressing these methodological and conceptual limitations—through multimodal imaging, longitudinal and experimental designs, diverse samples, and multi-level analyses—will clarify how attachment shapes the brain’s regulatory systems. The present study provides an initial step toward a neurophysiological oscillatory model of attachment-based emotion regulation, paving the way for translational research and individualized interventions.

## 5. Conclusions

Our study demonstrates that adult attachment orientations are reflected in distinct oscillatory dynamics within prefrontal brain networks during emotion regulation. Individuals higher in attachment anxiety exhibited reduced theta-band activity in the right dlPFC during cognitive reappraisal, suggesting diminished engagement of top-down cognitive control. In contrast, those higher in attachment avoidance showed increased beta-band activity in the left dlPFC during reappraisal, indicating a more rigid and cognitively distanced regulatory mode consistent with deactivating strategies (Engel & Fries, 2010; Klimesch, 2012; Vrtička & Vuilleumier, 2012).

These findings bridge attachment theory with neurophysiology, revealing that IWMs described by Bowlby (1969, 1982) have measurable neural correlates when individuals attempt to manage their emotions. The results align with and extend foundational and contemporary frameworks—from Bowlby and Ainsworth’s seminal ideas, through Mikulincer and Shaver’s (2007) hyperactivation/deactivation model, to the SoNeAt (White et al., 2023)—by providing concrete neurodynamic evidence of how attachment insecurity alters emotion regulation at the brain level. Moreover, these neural insights hold translational promise. Frontal theta and prefrontal beta oscillations may serve as potential biomarkers of regulatory capacity in insecurely attached individuals and as targets for interventions aimed at enhancing prefrontal control, such as neurofeedback or brain stimulation (Ros et al., 2013; Kang et al., 2014).

More broadly, this work underscores that attachment is not only a theory of social-emotional behavior but a biologically instantiated process: early relational experiences shape the wiring and oscillatory patterns of neural networks that sustain emotion regulation. By integrating attachment theory with cognitive neuroscience, we obtain a richer understanding of individual differences in emotion regulation—one that bridges interpersonal narratives and the brain’s electrical rhythms.

This integrative perspective invites future research to examine causality, developmental onset, and cultural influences, while encouraging clinicians to consider the neural as well as relational dimensions of emotion-regulation difficulties. Ultimately, fostering attachment security—through supportive relationships or targeted interventions—may yield benefits that are neurobiological as well as psychological, strengthening the brain’s capacity for adaptive emotional control. Prefrontal oscillatory dynamics thus offer a promising window into the interplay between attachment and emotion regulation, with implications for theory, clinical innovation, and wellbeing across the lifespan.

## Acknowledgements

We extend our gratitude to Milan Domic for his contribution to the creation and enhancement of the illustrations used in the emotional rating images and to Yolanda Schlumpf for generously sharing her emotion regulation task programming code. We also wish to thank Andrea Sánchez-Corzo, Martín Irani, Josefa Burgos, Catalina Carvallo, Leydi Granillo, Millaray Silva, Constanza Salazar, Josefa Silva, Loreto Fuentes, Franco Pizarro, Constanza Rore, and Lesly González for their assistance.

## 6. Funding

This research was financially supported by the Agencia Nacional de Investigación y Desarrollo (ANID) of Chile under the FONDECYT de Iniciación Grant number 11220009, FONDECYT regular 1250393, and FONDEQUIP EQM 230003.

## 7. Data availability

The data underlying this article will be shared upon reasonable request to the corresponding author.

---

**Supplementary Table S1.** Selected IAPS pictures per condition (Lang et al., 2005)

| N° | Experimental Conditions |  |  |  |
| --- | --- | --- | --- | --- |
|  | Nat-negative | Suppress | Reappraise | Nat-neutral |
| 1 | 2683 | 2095 | 2130 | 7009 |
| 2 | 2703 | 2205 | 2700 | 7011 |
| 3 | 2710 | 2900 | 2751 | 7012 |
| 4 | 2717 | 6211 | 2811 | 7018 |
| 5 | 3220 | 6231 | 3550 | 7042 |
| 6 | 3230 | 6250 | 6260 | 7045 |
| 7 | 3500 | 6312 | 6350 | 7061 |
| 8 | 6300 | 6560 | 6530 | 2190 |
| 9 | 6315 | 6563 | 6561 | 2215 |
| 10 | 6360 | 6838 | 6562 | 2359 |
| 11 | 6540 | 9041 | 6825 | 2499 |
| 12 | 6550 | 9250 | 6834 | 2518 |
| 13 | 6840 | 9421 | 9050 | 2521 |
| 14 | 9429 | 9427 | 9419 | 2593 |
| 15 | 9900 | 9435 | 9425 | 2594 |

Nat-negative: Natural condition containing negative valence pictures; Nat-neutral: Natural condition containing neutral valence pictures.

**Supplementary Table S2.** Descriptive statistics of the selected IAPS pictures per condition.

|  | Arousal |  |
| --- | --- | --- |
|  | Mean | SD |
| Nat-negative | 6.09 | 0.59 |
| Suppress | 5.76 | 0.81 |
| Reappraise | 5.83 | 0.81 |
| Nat-neutral | 3.48 | 0.48 |

Nat-negative: Natural condition containing negative valence pictures; Nat-neutral: Natural condition containing neutral valence pictures. SD = Standard Deviation.

**Supplementary Table S3.** Multiple comparison test for the arousal of IAPS pictures between conditions.

| Tukey's test | Mean Diff. | 95.00% CI of diff. | Summary | Adjusted p-value |
| --- | --- | --- | --- | --- |
| Natural-Neg vs. Suppress | 0.3240 | -0.3415 to 0.9895 | ns | 0.5736 |
| Natural-Neg vs. Reappraise | 0.2633 | -0.4022 to 0.9288 | ns | 0.7222 |
| **Natural-Neg vs. Natural-Neu** | **2.607** | **1.942 to 3.273** | **\*\*\*\*** | **<0.0001** |
| Suppress vs. Reappraise | -0.06067 | -0.7262 to 0.6048 | ns | 0.9950 |
| **Suppress vs. Natural-Neu** | **2.283** | **1.618 to 2.949** | **\*\*\*\*** | **<0.0001** |
| **Reappraise vs. Natural-Neu** | **2.344** | **1.679 to 3.009** | **\*\*\*\*** | **<0.0001** |

Natural-neg: Natural condition containing negative valence pictures; Natural-neu: Natural condition containing neutral valence pictures.; ns: not significant.

**Supplementary Table S4.**
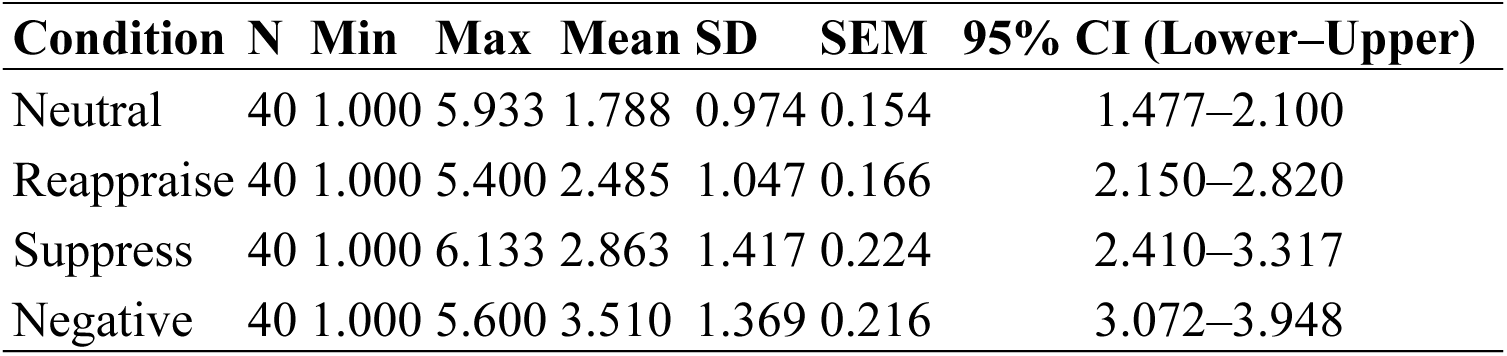
Descriptive statistics of self-reported arousal.

| Condition | N | Min | Max | Mean | SD | SEM | 95% CI (Lower–Upper) |
| --- | --- | --- | --- | --- | --- | --- | --- |
| Neutral | 40 | 1.000 | 5.933 | 1.788 | 0.974 | 0.154 | 1.477–2.100 |
| Reappraise | 40 | 1.000 | 5.400 | 2.485 | 1.047 | 0.166 | 2.150–2.820 |
| Suppress | 40 | 1.000 | 6.133 | 2.863 | 1.417 | 0.224 | 2.410–3.317 |
| Negative | 40 | 1.000 | 5.600 | 3.510 | 1.369 | 0.216 | 3.072–3.948 |

**Supplementary Table S5.** Shapiro–Wilk test of normality of self-reported arousal Condition W P value.

| Condition | W | P value |
| --- | --- | --- |
| Neutral | 0.7127 | < 0.0001 |
| Reappraise | 0.9534 | 0.0990 |
| Suppress | 0.9317 | 0.0182 |
| Negative | 0.9423 | 0.0412 |

**Supplementary Table S6.**
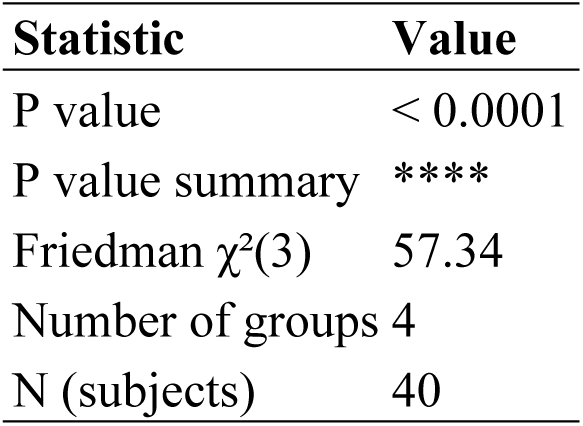
Friedman test summary of self-reported arousal Statistic Value.

**Supplementary Table S7.**
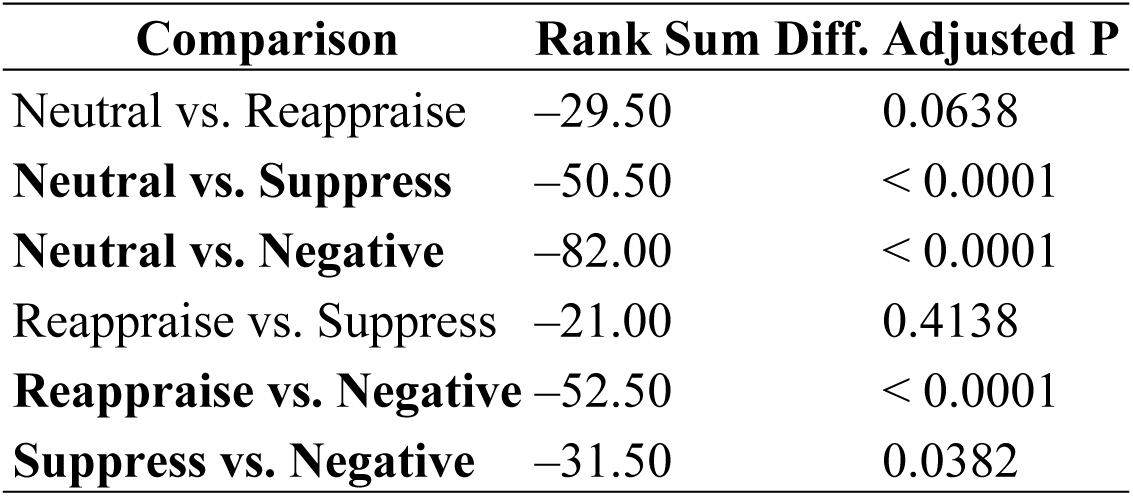
Dunn’s multiple comparisons test of self-reported arousal.

**Supplementary Table S8.**
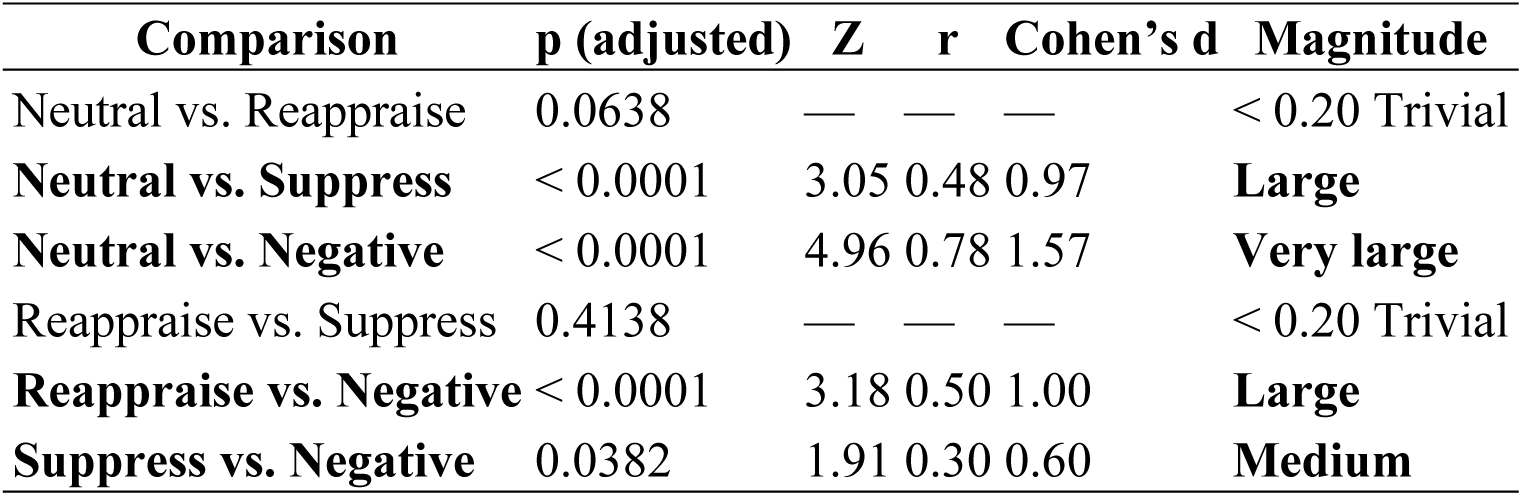
Post-hoc pairwise comparisons and effect sizes of self-reported arousal.

| Comparison | p (adjusted) | Z | r | Cohen's d | Magnitude |
| --- | --- | --- | --- | --- | --- |
| Neutral vs. Reappraise | 0.0638 | — | — | — | < 0.20 Trivial |
| **Neutral vs. Suppress** | < 0.0001 | 3.05 | 0.48 | 0.97 | **Large** |
| **Neutral vs. Negative** | < 0.0001 | 4.96 | 0.78 | 1.57 | **Very large** |
| Reappraise vs. Suppress | 0.4138 | — | — | — | < 0.20 Trivial |
| **Reappraise vs. Negative** | < 0.0001 | 3.18 | 0.50 | 1.00 | **Large** |
| **Suppress vs. Negative** | 0.0382 | 1.91 | 0.30 | 0.60 | **Medium** |

**Supplementary Table S9.** Theta band — Main effects and 2-way interactions (ML; N=960; DF=888)

| Effect | Estimate | SE | t | df | p | 95% CI |
| --- | --- | --- | --- | --- | --- | --- |
| (Intercept) | 0.028149 | 0.048934 | 0.57523 | 888 | 0.56528 | [-0.067892, 0.12419] |
| Condition_Reappraise | 0.024128 | 0.06593 | 0.36596 | 888 | 0.71448 | [-0.10527, 0.15353] |
| Condition_Suppress | 0.082297 | 0.068769 | 1.1967 | 888 | 0.23174 | [-0.052672, 0.21727] |
| ROI_L ACC | 0.03255 | 0.059136 | 0.55042 | 888 | 0.58217 | [-0.083513, 0.14861] |
| ROI_L dlPFC | 0.036987 | 0.059136 | 0.62545 | 888 | 0.53184 | [-0.079077, 0.15305] |
| ROI_L mPFC | 0.073217 | 0.059136 | 1.2381 | 888 | 0.216 | [-0.042846, 0.18928] |
| ROI_R ACC | 0.080577 | 0.059136 | 1.3626 | 888 | 0.17337 | [-0.035486, 0.19664] |
| ROI_R dlPFC | 0.10691 | 0.059136 | 1.8079 | 888 | 0.070962 | [-0.0091513, 0.22298] |
| ROI_R mPFC | 0.041956 | 0.059136 | 0.70947 | 888 | 0.47822 | [-0.074108, 0.15802] |
| ROI_R vlPFC / IFG | 0.017743 | 0.059136 | 0.30003 | 888 | 0.76422 | [-0.098321, 0.13381] |
| ANX_c | -0.018087 | 0.044787 | -0.40385 | 888 | 0.68642 | [-0.10599, 0.069813] |
| AVD_c | 0.015154 | 0.0413 | 0.36692 | 888 | 0.71377 | [-0.065903, 0.09621] |
| Condition_Reappraise:ROI_L ACC | 0.022228 | 0.082216 | 0.27037 | 888 | 0.78694 | [-0.13913, 0.18359] |
| Condition_Suppress:ROI_L ACC | -0.048065 | 0.082216 | -0.58462 | 888 | 0.55895 | [-0.20942, 0.11329] |
| Condition_Reappraise:ROI_L dlPFC | 0.042631 | 0.082216 | 0.51852 | 888 | 0.60422 | [-0.11873, 0.20399] |
| Condition_Suppress:ROI_L dlPFC | 0.019746 | 0.082216 | 0.24018 | 888 | 0.81025 | [-0.14161, 0.18111] |
| Condition_Reappraise:ROI_L mPFC | -0.004994 | 0.082216 | -0.06074 | 888 | 0.95158 | [-0.16635, 0.15637] |
| Condition_Suppress:ROI_L mPFC | -0.13449 | 0.082216 | -1.6359 | 888 | 0.10222 | [-0.29585, 0.026867] |
| Condition_Reappraise:ROI_R ACC | -0.027419 | 0.082216 | -0.3335 | 888 | 0.73883 | [-0.18878, 0.13394] |
| Condition_Suppress:ROI_R ACC | -0.080605 | 0.082216 | -0.98041 | 888 | 0.32715 | [-0.24196, 0.080755] |
| Condition_Reappraise:ROI_R dlPFC | -0.029588 | 0.082216 | -0.35988 | 888 | 0.71902 | [-0.19095, 0.13177] |
| Condition_Suppress:ROI_R dlPFC | -0.13908 | 0.082216 | -1.6916 | 888 | 0.09107 | [-0.30044, 0.022282] |
| Condition_Reappraise:ROI_R mPFC | 0.059035 | 0.082216 | 0.71805 | 888 | 0.47292 | [-0.10232, 0.22039] |
| Condition_Suppress:ROI_R mPFC | -0.065274 | 0.082216 | -0.79394 | 888 | 0.42744 | [-0.22663, 0.096085] |
| Condition_Reappraise:ROI_R vlPFC / IFG | 0.0053231 | 0.082216 | 0.06475 | 888 | 0.94839 | [-0.15604, 0.16668] |
| Condition_Suppress:ROI_R vlPFC / IFG | -0.058369 | 0.082216 | -0.70995 | 888 | 0.47792 | [-0.21973, 0.10299] |
| Condition_Reappraise:ANX_c | 0.030524 | 0.060342 | 0.50586 | 888 | 0.61308 | [-0.087905, 0.14895] |
| Condition_Suppress:ANX_c | -0.026955 | 0.06294 | -0.42826 | 888 | 0.66856 | [-0.15048, 0.096574] |
| ROI_L ACC:ANX_c | -0.0197 | 0.054124 | -0.36398 | 888 | 0.71596 | [-0.12593, 0.086526] |
| ROI_L dlPFC:ANX_c | -0.056714 | 0.054124 | -1.0478 | 888 | 0.29499 | [-0.16294, 0.049512] |
| ROI_L mPFC:ANX_c | 0.019152 | 0.054124 | 0.35385 | 888 | 0.72354 | [-0.087074, 0.12538] |
| ROI_R ACC:ANX_c | -0.034821 | 0.054124 | -0.64336 | 888 | 0.52016 | [-0.14105, 0.071405] |
| ROI_R dlPFC:ANX_c | 0.040166 | 0.054124 | 0.7421 | 888 | 0.45822 | [-0.06606, 0.14639] |
| ROI_R mPFC:ANX_c | 0.011717 | 0.054124 | 0.21648 | 888 | 0.82866 | [-0.094509, 0.11794] |
| ROI_R vlPFC / IFG:ANX_c | 0.060895 | 0.054124 | 1.1251 | 888 | 0.26085 | [-0.045331, 0.16712] |
| Condition_Reappraise:AVD_c | 0.043305 | 0.055644 | 0.77825 | 888 | 0.43663 | [-0.065904, 0.15251] |
| Condition_Suppress:AVD_c | -0.062006 | 0.05804 | -1.0683 | 888 | 0.28566 | [-0.17592, 0.051906] |
| ROI_L ACC:AVD_c | 0.00094817 | 0.04991 | 0.018997 | 888 | 0.98485 | [-0.097007, 0.098904] |
| ROI_L dlPFC:AVD_c | -0.078228 | 0.04991 | -1.5674 | 888 | 0.11738 | [-0.17618, 0.019727] |
| ROI_L mPFC:AVD_c | -0.018024 | 0.04991 | -0.36112 | 888 | 0.71809 | [-0.11598, 0.079932] |
| ROI_R ACC:AVD_c | 0.03331 | 0.04991 | 0.66739 | 888 | 0.5047 | [-0.064646, 0.13127] |
| ROI_R dlPFC:AVD_c | -0.022212 | 0.04991 | -0.44505 | 888 | 0.6564 | [-0.12017, 0.075743] |
| ROI_R mPFC:AVD_c | -0.0083151 | 0.04991 | -0.1666 | 888 | 0.86772 | [-0.10627, 0.089641] |
| ROI_R vlPFC / IFG:AVD_c | -0.0050048 | 0.04991 | -0.10028 | 888 | 0.92015 | [-0.10296, 0.092951] |

**Supplementary Table S10.** Theta band — 3-way interactions.

| Effect | Estimate | SE | t | df | p | 95% CI |
| --- | --- | --- | --- | --- | --- | --- |
| Condition_Reappraise:ROI_L ACC:ANX_c | -0.098859 | 0.075247 | -1.3138 | 888 | 0.18926 | [-0.24654, 0.048825] |
| Condition_Suppress:ROI_L ACC:ANX_c | -0.028603 | 0.075247 | -0.38012 | 888 | 0.70395 | [-0.17629, 0.11908] |
| Condition_Reappraise:ROI_L dlPFC:ANX_c | 0.015424 | 0.075247 | 0.20498 | 888 | 0.83763 | [-0.13226, 0.16311] |
| Condition_Suppress:ROI_L<br>dlPFC:ANX_c | 0.063355 | 0.075247 | 0.84196 | 888 | 0.40004 | [-0.084328,<br>0.21104] |
| Condition_Reappraise:ROI_L<br>mPFC:ANX_c | -0.10936 | 0.075247 | -1.4534 | 888 | 0.14647 | [-0.25705,<br>0.038321] |
| Condition_Suppress:ROI_L<br>mPFC:ANX_c | -0.085503 | 0.075247 | -1.1363 | 888 | 0.25614 | [-0.23319,<br>0.06218] |
| Condition_Reappraise:ROI_R<br>ACC:ANX_c | -0.056179 | 0.075247 | -0.7466 | 888 | 0.4555 | [-0.20386,<br>0.091504] |
| Condition_Suppress:ROI_R<br>ACC:ANX_c | -0.0049498 | 0.075247 | -0.06578 | 888 | 0.94757 | [-0.15263,<br>0.14273] |
| **Condition_Reappraise:ROI_R**<br>**dlPFC:ANX_c** | **-0.18656** | **0.075247** | **-2.4793** | **888** | **0.013347** | **[-0.33425, -**<br>**0.03888]** |
| Condition_Suppress:ROI_R<br>dlPFC:ANX_c | -0.013956 | 0.075247 | -0.18547 | 888 | 0.8529 | [-0.16164,<br>0.13373] |
| Condition_Reappraise:ROI_R<br>mPFC:ANX_c | -0.095767 | 0.075247 | -1.2727 | 888 | 0.20346 | [-0.24345,<br>0.051916] |
| Condition_Suppress:ROI_R<br>mPFC:ANX_c | -0.075781 | 0.075247 | -1.0071 | 888 | 0.31416 | [-0.22346,<br>0.071902] |
| Condition_Reappraise:ROI_R<br>vlPFC / IFG:ANX_c | -0.02043 | 0.075247 | -0.27151 | 888 | 0.78606 | [-0.16811,<br>0.12725] |
| Condition_Suppress:ROI_R<br>vlPFC / IFG:ANX_c | -0.069451 | 0.075247 | -0.92297 | 888 | 0.35627 | [-0.21713,<br>0.078232] |
| Condition_Reappraise:ROI_L<br>ACC:AVD_c | -0.023202 | 0.069389 | -0.33438 | 888 | 0.73818 | [-0.15939,<br>0.11298] |
| Condition_Suppress:ROI_L<br>ACC:AVD_c | 0.085975 | 0.069389 | 1.239 | 888 | 0.21566 | [-0.05021,<br>0.22216] |
| Condition_Reappraise:ROI_L<br>dlPFC:AVD_c | 0.089594 | 0.069389 | 1.2912 | 888 | 0.19697 | [-0.046591,<br>0.22578] |
| Condition_Suppress:ROI_L<br>dlPFC:AVD_c | 0.10655 | 0.069389 | 1.5356 | 888 | 0.125 | [-0.029633,<br>0.24274] |
| Condition_Reappraise:ROI_L<br>mPFC:AVD_c | -0.018569 | 0.069389 | -0.2676 | 888 | 0.78907 | [-0.15475,<br>0.11762] |
| Condition_Suppress:ROI_L<br>mPFC:AVD_c | 0.13878 | 0.069389 | 2.0000 | 888 | **0.045801** | [0.002595,<br>0.27496] |
| Condition_Reappraise:ROI_R<br>ACC:AVD_c | -0.097556 | 0.069389 | -1.4059 | 888 | 0.1601 | [-0.23374,<br>0.038629] |
| Condition_Suppress:ROI_R<br>ACC:AVD_c | 0.022813 | 0.069389 | 0.32877 | 888 | 0.74241 | [-0.11337,<br>0.159] |
| Condition_Reappraise:ROI_R<br>dlPFC:AVD_c | -0.059808 | 0.069389 | -0.86193 | 888 | 0.38896 | [-0.19599,<br>0.076377] |
| Condition_Suppress:ROI_R<br>dlPFC:AVD_c | 0.037291 | 0.069389 | 0.53742 | 888 | 0.59111 | [-0.098894,<br>0.17348] |
| Condition_Reappraise:ROI_R<br>mPFC:AVD_c | -0.077131 | 0.069389 | -1.1116 | 888 | 0.26662 | [-0.21332,<br>0.059054] |
| Condition_Suppress:ROI_R<br>mPFC:AVD_c | 0.13961 | 0.069389 | 2.012 | 888 | **0.044526** | [0.003422,<br>0.27579] |
| Condition_Reappraise:ROI_R<br>vlPFC / IFG:AVD_c | -0.0839 | 0.069389 | -1.2091 | 888 | 0.22694 | [-0.22008,<br>0.052285] |
| Condition_Suppress:ROI_R<br>vlPFC / IFG:AVD_c | 0.079416 | 0.069389 | 1.1445 | 888 | 0.25272 | [-0.056769,<br>0.2156] |

**Supplementary Table S11.**
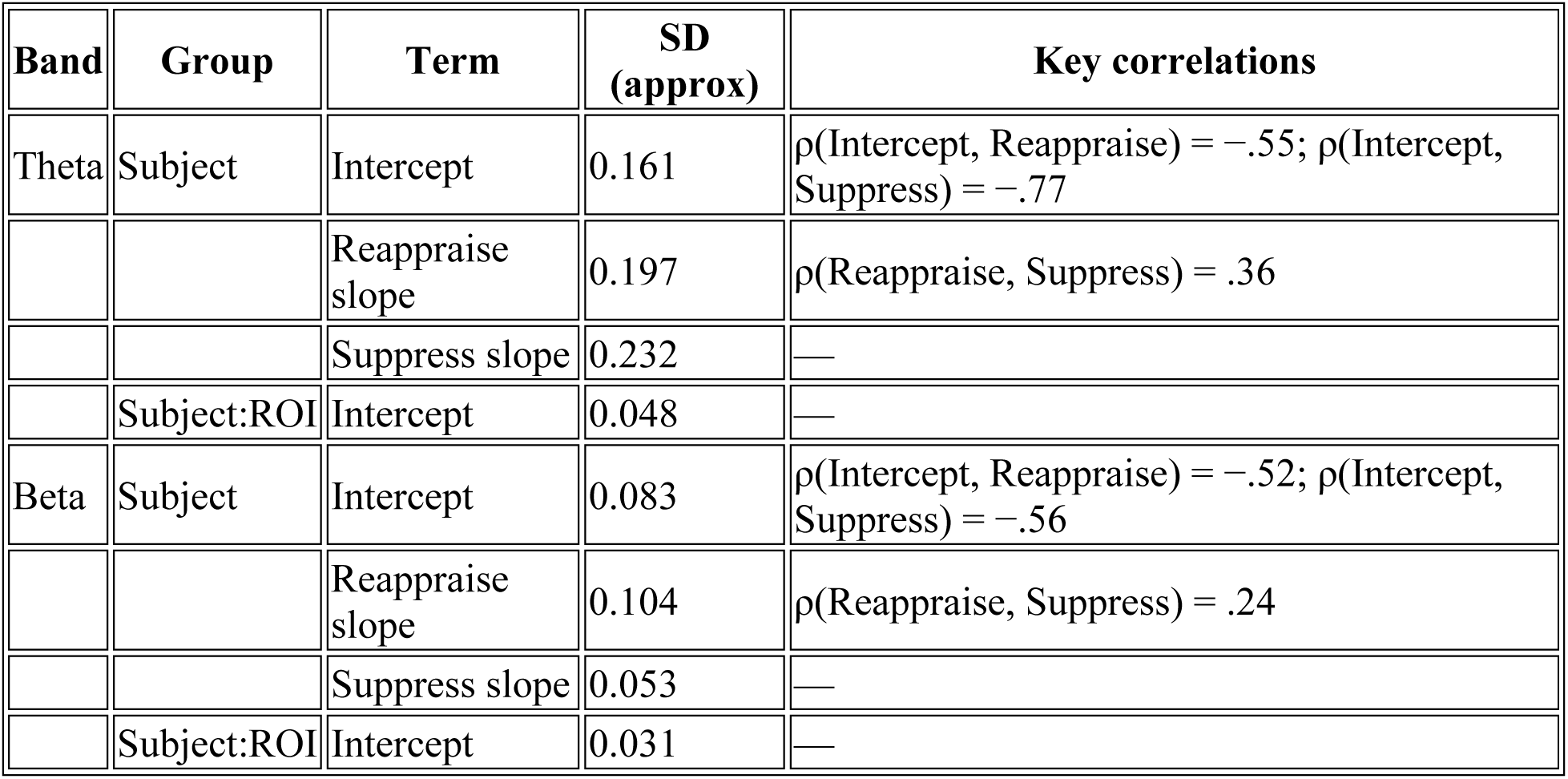
Random-effects (SD/correlations; 95% CIs in text)

**Supplementary Table S12.**
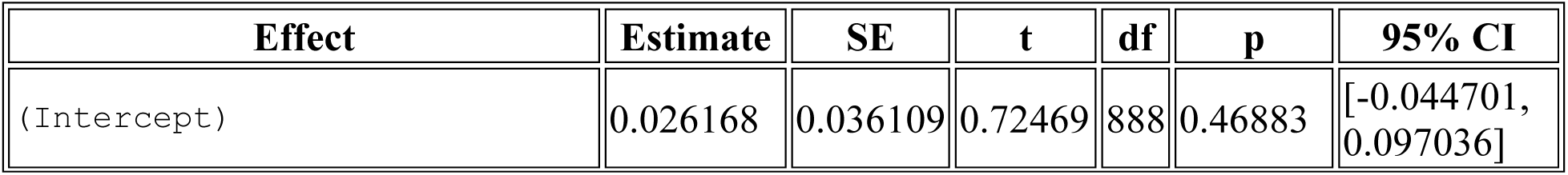
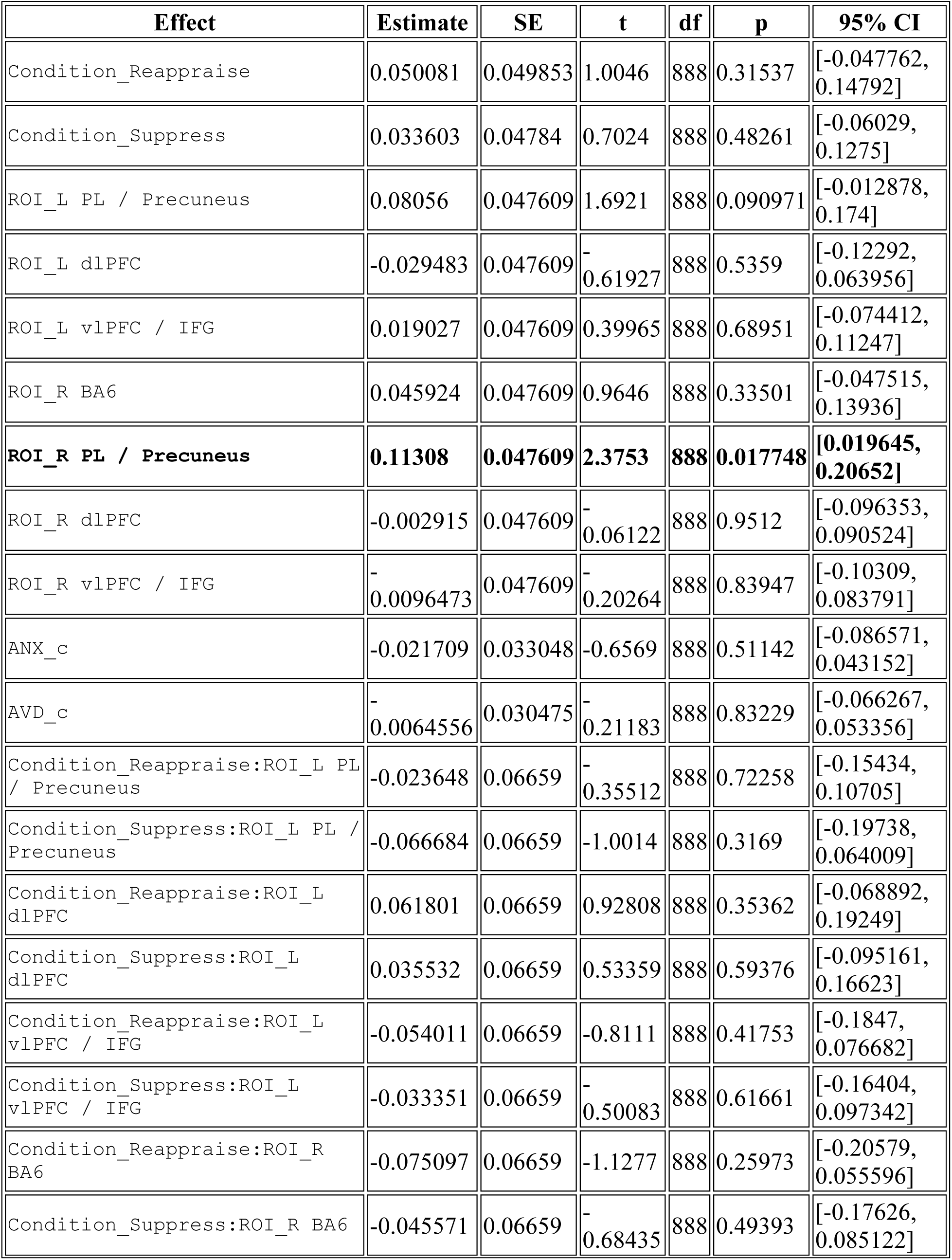
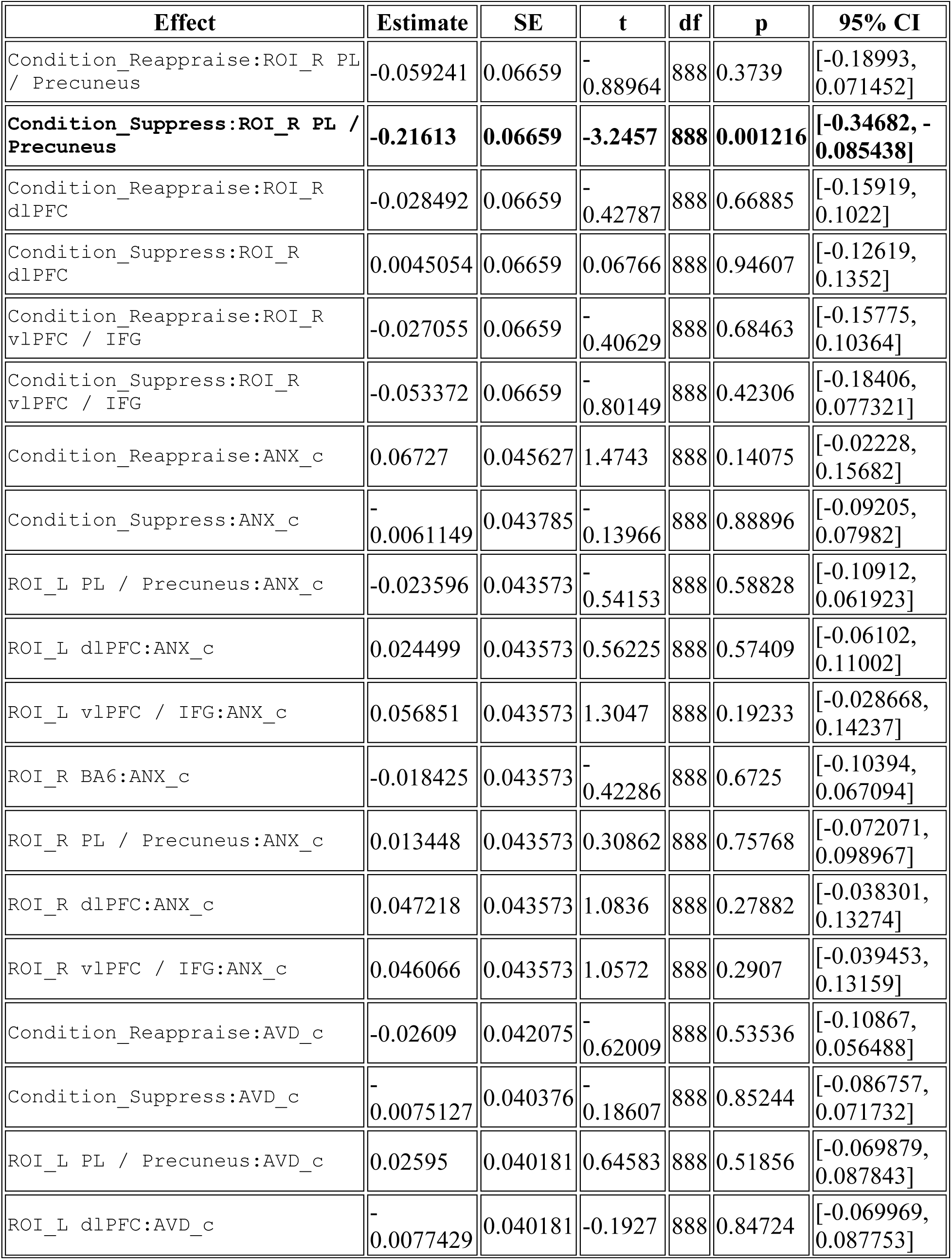
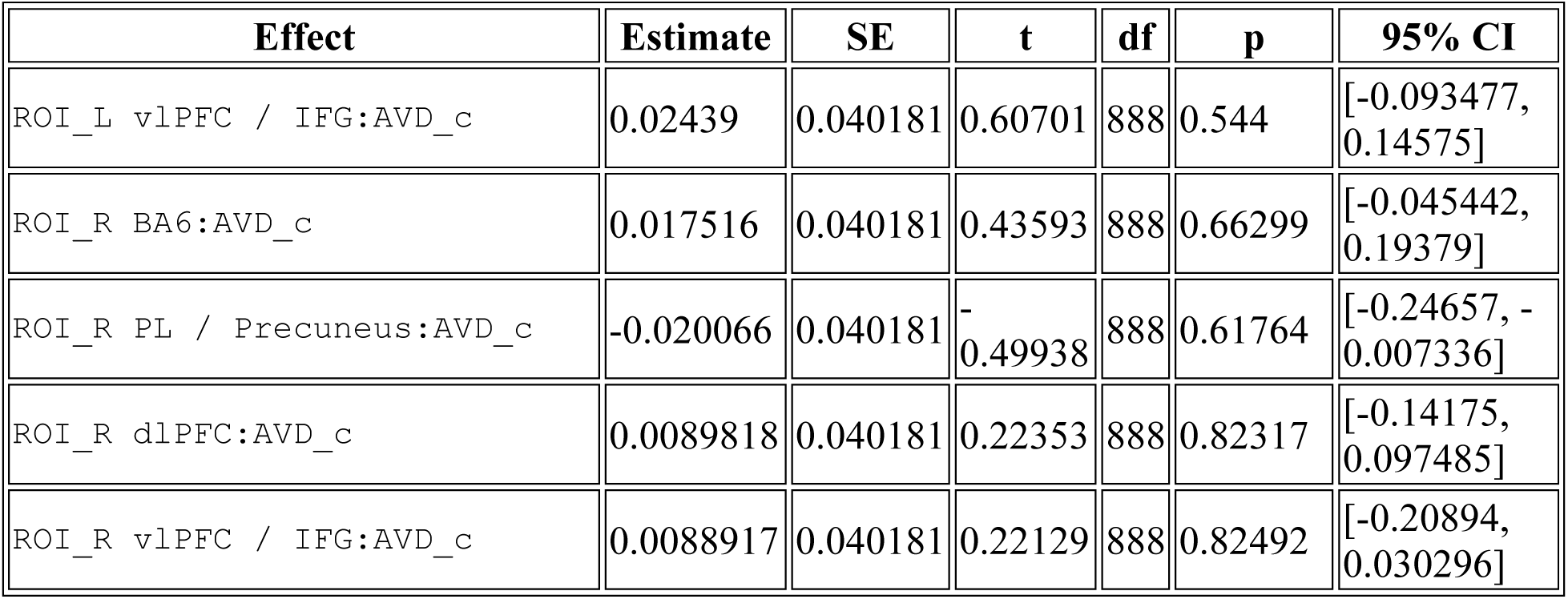
Beta band — Main effects and 2-way interactions (ML; N=960; DF=888)

**Supplementary Table S13.**
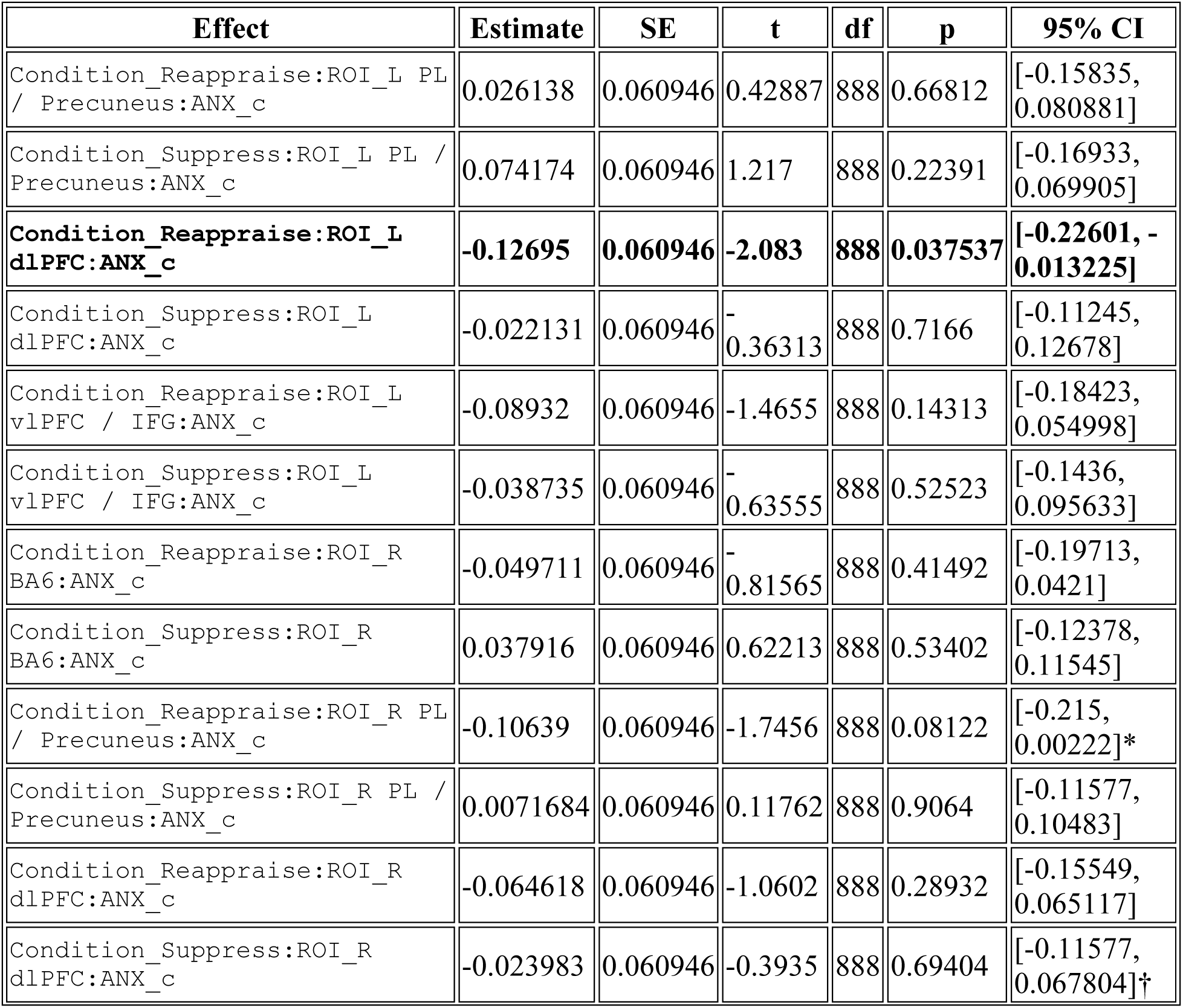
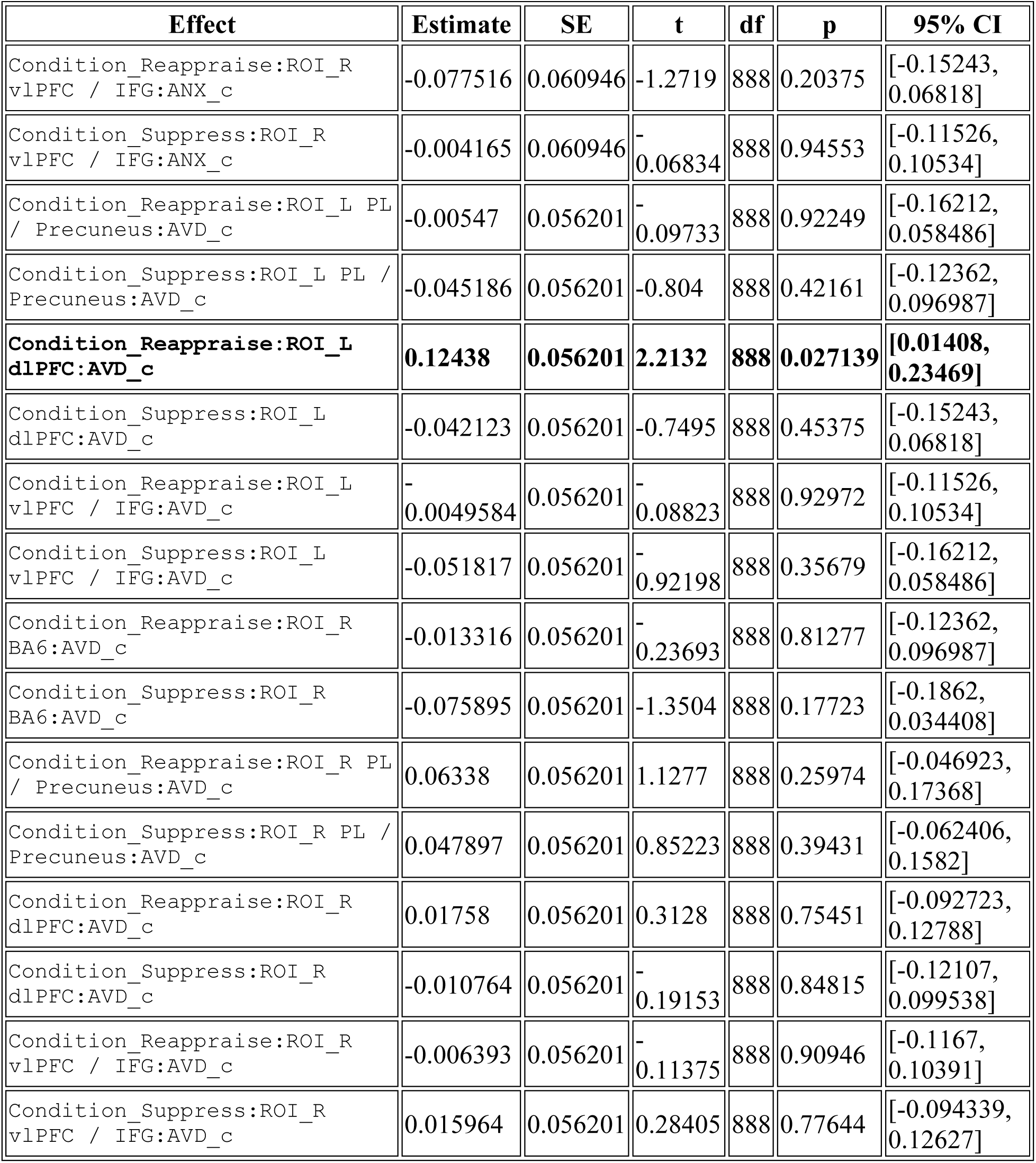
Beta band — 3-way interactions.

## Supplementary Figures

**Supplementary Figure S1.**
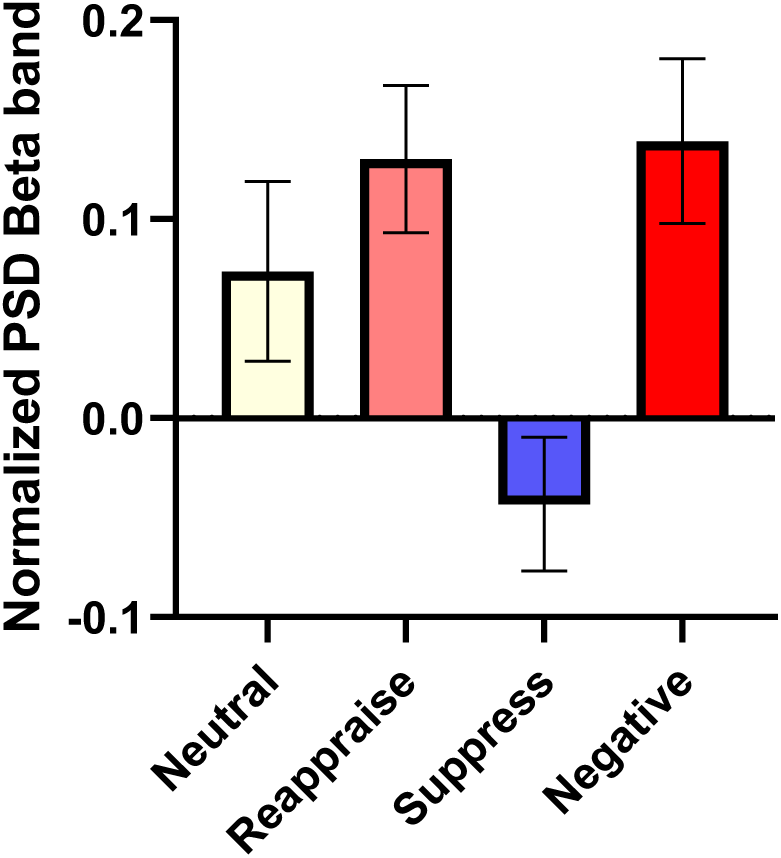
Normalized beta-band (15–30 Hz) power in the right parietal/precuneus (R PL/Precuneus) region across emotion regulation conditions. Bars represent mean normalized power spectral density (PSD) values (A–B)/(A+B) relative to baseline for Neutral, Reappraise, Suppress, and Negative conditions (N = 40). Beta activity showed a clear decrease during Suppress (M = –0.04, SD = 0.21), in contrast to overall positive modulation during Neutral (M = 0.07, SD = 0.29), Reappraise (M = 0.13, SD = 0.23), and Negative (M = 0.14, SD = 0.26) conditions. Error bars denote ±1 SEM. This pattern illustrates the task-related beta desynchronization specific to suppression, consistent with reduced parietal readiness and visuo-attentional engagement during expressive inhibition.

## Notes

### Competing Interest Statement

The authors have declared no competing interest.

